# Integrating cell-state and enhancer constraints discriminates models of PU.1 regulation during hematopoietic differentiation

**DOI:** 10.64898/2026.10.01.756127

**Authors:** Yujia Zhang, Oleg Igoshin

**Affiliations:** Department of Bioengineering, Rice University, Houston, Texas, USA; Departments of Chemistry and of Biosciences, Center for Theoretical Biological Physics, and Rice Synthetic Biology Institute, Rice University, Houston, Texas, USA

## Abstract

Distinct PU.1 levels are associated with progenitor, B-cell, and macrophage states during hematopoietic differentiation. A previous dynamical-systems model attributed these levels to multistability generated by PU.1 autoregulation coupled to feedback between PU.1 abundance and cell-cycle progression, but its predictions were not quantitatively evaluated against the motivating measurements. Here, we test this mechanism using reported measurements of PU.1 abundance, its synthesis rate, and cell-cycle duration, and develop progressively more mechanistic alternatives. The original growth-feedback model neither represents the two reported progenitor groups nor reproduces the nonmonotonic relationship between PU.1 abundance and cell-cycle length. Even with experimentally measured cell-type-specific growth rates, PU.1 autoregulation alone does not reproduce the observed PU.1 abundance and synthesis rates across cell states. Reproducing the cell-state measurements requires an additional cell-state-dependent input to PU.1 production. Guided by enhancer regulation, we model positive feedback between PU.1 and C/EBP*α* and constrain it with reporter measurements, but this architecture cannot jointly reproduce the reporter and cell-state datasets. Adding RUNX1 permits data-consistent stable states when high effective cooperativity in PU.1-dependent C/EBP*α* activation is allowed. Thus, combined dynamical and cis-regulatory constraints support additional regulation of PU.1 production beyond autoregulation, cell-type-specific dilution, and C/EBP*α* feedback. The RUNX1-extended architecture provides one biologically grounded realization and generates experimentally testable hypotheses for differentiation networks.

## 1 Introduction

Hematopoietic stem cells (HSCs) sustain blood production by generating progenitors that progressively acquire restricted lineage potential [1, 2]. Although classical roadmaps describe an early bifurcation between common myeloid and common lymphoid progenitors [3, 4], single-cell and lineage-tracing studies indicate that hematopoietic differentiation is more heterogeneous than a simple binary tree [5–7]. A central challenge is therefore to determine how regulatory networks generate and stabilize lineage-associated molecular states.

The transcription factor PU.1 is a key regulator of this process [8–11]. PU.1 abundance increases as HSCs enter multipotent progenitor states [5, 6, 12] and subsequently differs between mature lineages, with higher levels generally associated with macrophages and lower levels with B cells [4, 5, 13]. These graded levels can influence lineage-associated transcriptional programs, making PU.1 a useful system for examining how quantitative differences in a master regulator are established and maintained.

Several mechanisms regulate PU.1 abundance. PU.1 activates its own transcription through promoter and enhancer elements [13–15], while C/EBP*α*, RUNX1, GATA-1, and other regulators also act at the PU.1 locus [16–18]. PU.1 can additionally slow cell-cycle progression by repressing cell-cycle activators or their regulators [19–21]. Because cell division dilutes intracellular proteins, changes in cell-cycle length could, in turn, alter PU.1 concentration. PU.1 levels may therefore emerge from the combined effects of transcriptional regulation, protein turnover, and cell division.

A previous experimental and modeling study proposed that PU.1 autoregulation and growth feedback jointly generate lineage-associated PU.1 states [5]. In that autoregulation- and-growth-feedback (AR&GF) model, increasing PU.1 slows cell division, reduced dilution permits further PU.1 accumulation, and the resulting positive feedback produces three stable steady states assigned to B cells, progenitors, and macrophages. This work provided an influential dynamical-systems interpretation of PU.1 regulation and reported single-cell measurements of PU.1-GFP abundance, PU.1 synthesis rate, and cell-cycle length in B cells, macrophages, and two operationally defined progenitor groups. The study referred to these groups as pro-low and pro-high based on their relative PU.1 synthesis rates and proposed that they were associated with different lineage outcomes. However, their identities and differentiation trajectories were not directly established by immunophenotyping.

The AR&GF model, however, was developed primarily to explain qualitative trends. It predicts one progenitor attractor rather than the two groups suggested by the measurements, and its assumption that cell-cycle length increases monotonically with PU.1 is not supported across the reported cell types. More broadly, it remains unclear which regulatory features are minimally required to reproduce both the cell-state measurements and independent perturbation data on PU.1 transcription. Addressing this question requires treating PU.1 abundance, synthesis-rate, and cell-cycle measurements as constraints on the steady states and removal rates, while using enhancer-reporter perturbations to constrain the PU.1 production function. This integrated approach can distinguish models that merely generate multistability from those that are quantitatively consistent with multiple data types.

Here, we use a sequence of phenomenological and biophysical models to identify the regulatory features required within the tested model class to satisfy these complementary cell-state and transcriptional constraints. We first quantify the disagreement between the AR&GF model and the PU.1 abundance, synthesis-rate, and cell-cycle data. We then replace growth feedback with measured, cell-type-specific dilution rates and test whether PU.1 autoregulation alone is sufficient. After identifying the need for cell-type-dependent regulation of PU.1 production, we evaluate positive feedback with C/EBP*α* using a biophysical model constrained by PU.1 reporter data and finally test the effect of RUNX1 input. This model-discrimination strategy identifies C/EBP*α* and RUNX1 as a biologically plausible implementation of the additional regulatory input required within the tested model class, while generating specific hypotheses for experimental evaluation.

## 2 Results

### 2.1 The published autoregulation-and-growth-feedback model is quantitatively inconsistent with the reported measurements

We first asked whether the previously proposed autoregulation-and-growth-feedback (AR&GF) model quantitatively reproduces the measurements used to motivate it [5]. In this model, PU.1 activates its own production, while increasing PU.1 slows cell division and thereby reduces dilution (Figure 1A). The two positive-feedback loops generate three stable steady states that were assigned to B cells, progenitor cells, and macrophages (Figure 1B). To compare these predictions with the reported PU.1-GFP abundance, PU.1 synthesis rate, and cell-cycle length, we expressed both the model outputs and experimental measurements relative to the macrophage values (Methods 5.1 and 5.2). Because the macrophage state defines the normalization, its dimensionless PU.1 level, synthesis rate, and cell-cycle length are fixed at one and do not constitute independent predictions of the model.

**Figure 1.**
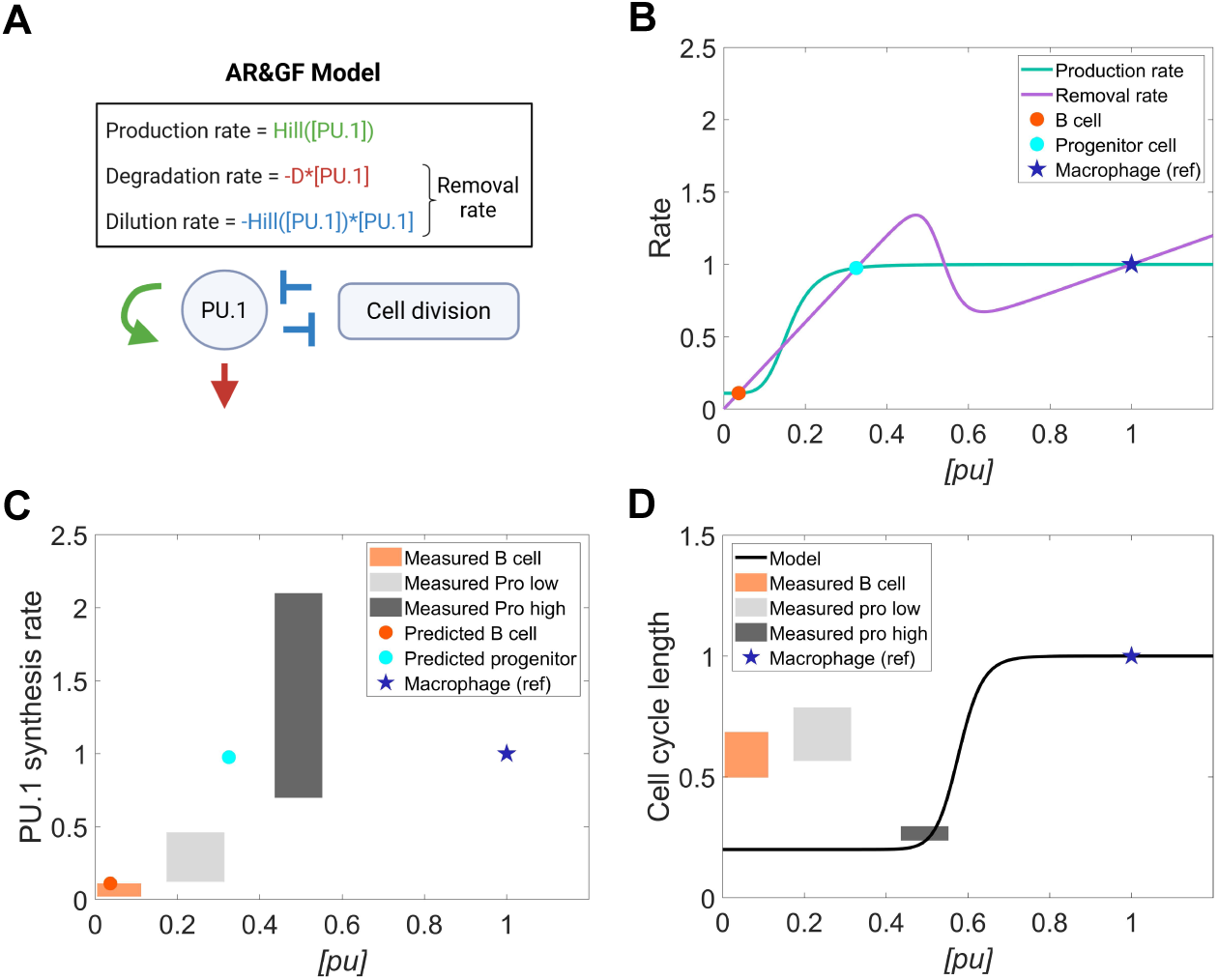
The “AR&GF” model is quantitatively inconsistent with the measured PU.1 abundance, PU.1 synthesis rate, and cell-cycle lengths. **(A)** The architecture of the “AR&GF model” is shown. PU.1 is produced through a self-activating phenomenological function (green arrow) and removed through natural degradation (red arrow) and cell division (blue arrows). The dilution rate constant decreases with PU.1 abundance in a Hill manner. **(B)** Predictions of the “AR&GF model” are visualized as intersections of the PU.1-dependent production and removal functions. Of the five intersections (steady states), three are stable and were assigned to distinct cell states during HSC differentiation. The plotted dimensionless model was obtained by normalizing all model equations by the macrophage PU.1 level (Methods 5.2). Lowercase italic “[pu]” represents the dimensionless PU.1 concentration. **(C)** Model-predicted dimensionless PU.1 levels and production rates (circle markers) are compared with the joint experimental uncertainty ranges (colored patches). In subsequent figures, the colored patches denote the same joint measurement ranges and are not repeated in the legends. For each cell type, the measured PU.1 level and PU.1 synthesis rates are normalized by the median of the macrophage measurements. **(D)** The model-imposed relationship between cell-cycle length and PU.1 level is compared with the measurements. The model cell-cycle length (black line) is derived from the dilution rate constant in (A), assuming exponential cell growth. The measured cell-cycle length is normalized to the mean macrophage cycle length. Details of uncertainty propagation are provided in Supplementary Section 1. Because both the model and data were normalized, the macrophage state has a dimensionless PU.1 level, PU.1 production rate, and cell-cycle length of 1, and necessarily agrees with the data. The macrophage is therefore a reference state (“ref”, blue star marker). All model parameters and data were taken from [5].

The first discrepancy concerns the number of progenitor states. The AR&GF model contains a single intermediate stable state (Figure 1B), whereas the source study reported two operationally defined progenitor groups with distinct PU.1-GFP levels, synthesis rates, and cell-cycle lengths [5]. Retaining the terminology of [5], we refer to these groups as pro-low and pro-high. The original study implied that pro-low cells develop toward the B-cell lineage and pro-high cells toward the macrophage lineage, but these trajectories were not established by direct immunophenotyping.

The model does not reproduce the measured PU.1 abundance and synthesis rates across these groups. Its low-PU.1 stable state falls within the reported B-cell range for both quantities (Figure 1C). In contrast, the single intermediate state lies outside the joint ranges measured for both pro-low and pro-high. Thus, after normalization to the macrophage reference state, the AR&GF model reproduces the relative PU.1 abundance and synthesis rate measured in B cells but does not quantitatively represent either progenitor group.

The growth-feedback function is also inconsistent with the relationship between PU.1 level and cell-cycle length across the reported cell types. In the AR&GF model, cell-cycle length increases monotonically with PU.1 (Figure 1D), so higher PU.1 necessarily produces slower division and less dilution. The measurements do not follow this ordering. In particular, pro-high cells have the highest PU.1 level but the shortest reported cell-cycle length, whereas pro-low cells and B cells cycle more slowly despite having lower PU.1 levels. Consequently, the model-predicted curve does not overlap the measured ranges for B cells or pro-low cells. The feedback proposed from the concurrent increases in PU.1 and cell-cycle length during macrophage maturation, therefore, does not generalize across all four measured groups.

Together, these comparisons identify two quantitative limitations of the published mechanism: it provides only one progenitor attractor, and it imposes a universal monotonic relationship between PU.1 and cell-cycle length that is not supported across cell types. We therefore asked whether the core idea of PU.1 autoregulation could be retained while replacing growth feedback with cell-type-specific dilution rates directly constrained by measured cell-cycle lengths.

### 2.2 Measured cell-type-specific dilution is insufficient for a model based on PU.1 autoregulation alone

The quantitative disagreement with the AR&GF model could arise from its assumption that PU.1 determines cell-cycle length through a common growth-feedback function. We therefore tested whether PU.1 autoregulation could explain the measured states if growth feedback was removed and the dilution rate of each cell type was instead constrained directly by its reported mean cell-cycle length. Notably, for pro-high, we used the non-mean value (approximately the 75th percentile) as the cell-cycle length constraint (Methods 5.3). In this autoregulation-and-discrete-growth-rate (AR&DGR) model, PU.1 is produced through the same self-activating Hill function in every cell type, degraded at a common constant rate, and diluted at a cell-type-specific constant rate (Figure 2A; Methods 5.4). Thus, the production mechanism and its parameters are shared across cell types, whereas differences in removal arise only from the measured cell-cycle lengths.

**Figure 2.**
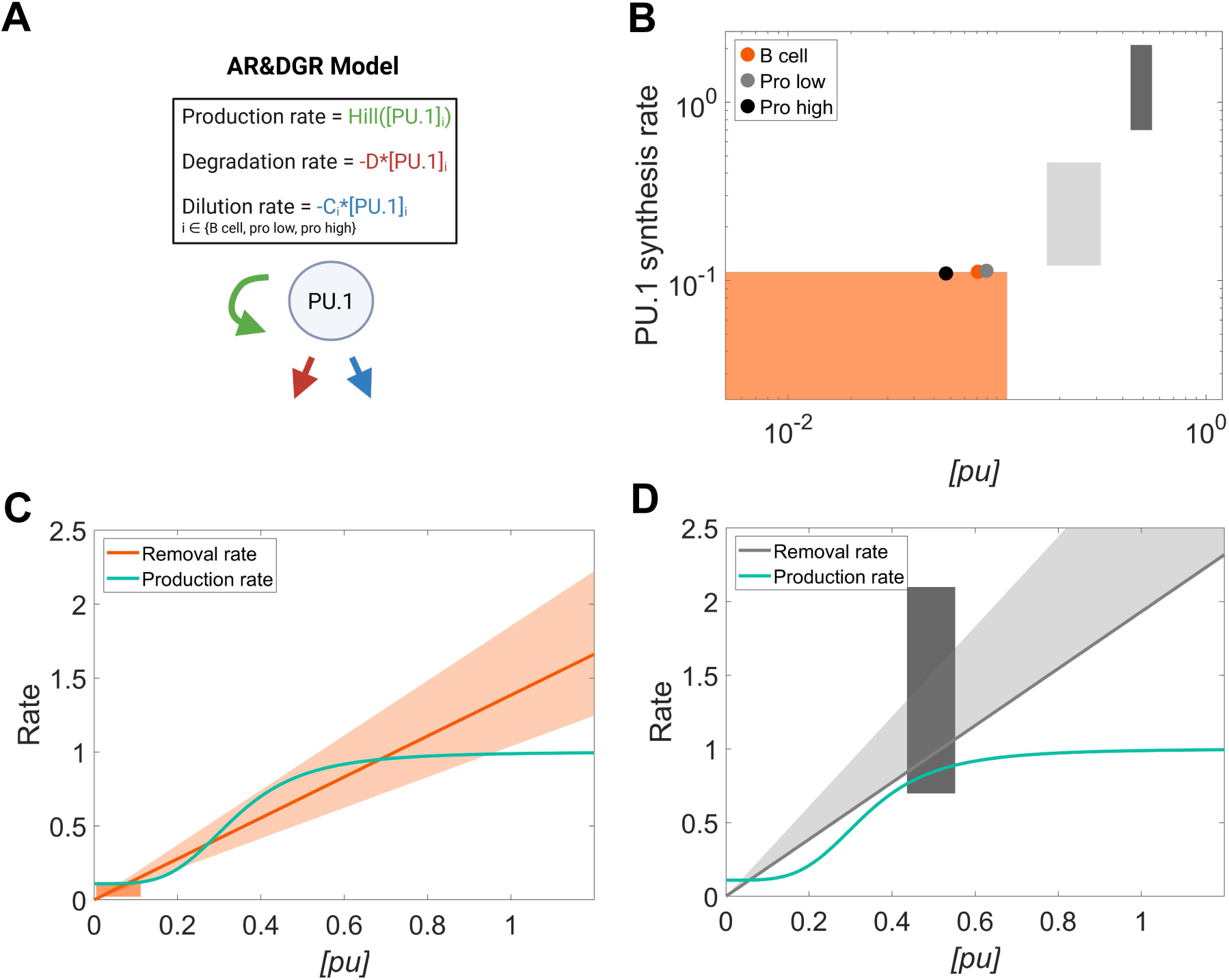
A phenomenological model with cell-type-specific growth rates and PU.1 autoregulation failed to reproduce the observed PU.1 abundance and PU.1 synthesis rates of B cells, pro-low cells, and pro-high cells. **(A)** The architecture of the “AR&DGR model” is shown. PU.1 is produced with a self-activating Hill function (green arrow), degraded at a constant rate (red arrow), and diluted at a constant rate (blue arrow). The dilution rate constant is constrained by the measured cell-cycle length [5]. In contrast to the “AR&GF model” in Figure 1A, each cell type “i” is represented by a separate equation with its own removal rate (Methods 5.3). **(B)** Fitted PU.1 concentrations and production rates (circle markers) are compared with the experimental ranges (colored patches). **(C)** The fitted phenomenological production function is plotted against the protein removal function (constrained by cell-cycle length) of the B cell. **(D)** The same production function is plotted against the removal function for pro-high cells. In both (C) and (D), the shaded region around the removal rate represents uncertainty propagated from experimental cell-cycle length [5].

We fitted the AR&DGR model simultaneously to the normalized PU.1 abundance and synthesis-rate ranges for B cells, pro-low cells, and pro-high cells (Figure 2B). As in the preceding analysis, macrophages define the reference state and are not treated as an independent prediction. When a fitted equation produced multiple stable states for a cell type, we evaluated the stable state closest to that cell type’s measured PU.1 abundance range. Under these conditions, the model identifies a B-cell state that falls within the reported ranges for both PU.1 abundance and synthesis rate. In contrast, the fitted states for pro-low and pro-high lie outside their corresponding joint measurement ranges. Incorporating the observed differences in cell-cycle length is therefore insufficient to make PU.1 autoregulation quantitatively consistent with the two progenitor groups.

The model failure can be understood by comparing its production and removal functions. At steady state, the PU.1 production rate must equal the total removal rate, which comprises natural degradation and dilution due to cell division. Because cell-cycle lengths differ among the measured groups, each cell type has a distinct linear removal function. The AR&DGR model, however, requires a single shared autoregulatory Hill function to intersect each removal function at a PU.1 level and a production rate that both fall within the corresponding experimental ranges.

No shared production function satisfies these constraints across all three non-reference groups. For B cells, the fitted production function intersects the removal function within the measured PU.1 range (Figure 2C). The same production function does not generate an acceptable pro-high state when paired with the pro-high removal function (Figure 2D), and analogous comparisons show the mismatch for pro-low (Supplementary Figure S1). This limitation is not simply a failure to produce enough steady states. Rather, the measured PU.1 levels and cell-type-specific removal rates imply different required production rates that cannot all be generated by a single PU.1-dependent Hill function with shared parameters.

These results distinguish the effects of PU.1 removal from those of PU.1 production. Replacing growth feedback with measured cell-type-specific dilution eliminates the unsupported assumption of a universal PU.1-to-cell-cycle relationship, but it does not resolve the quantitative mismatch. The remaining failure indicates that PU.1 production must depend on an additional cell-state-associated input beyond PU.1 autoregulation. We next introduce the minimal such input without initially assigning it a specific molecular identity.

### 2.3 A cell-state-dependent input to PU.1 production reproduces the measured states

The failure of the AR&DGR model indicated that measured differences in cell-cycle-dependent removal were not sufficient to explain the reported PU.1 states. We therefore asked whether an additional cell-state-dependent contribution to PU.1 production could account for the remaining differences. To test this possibility without initially assigning a molecular mechanism, we multiplied the shared PU.1 autoregulatory Hill function by a factor, X, that was fitted independently for B cells, pro-low cells, and pro-high cells (Figure 3A). In this “cell-type-specific-autoregulation model”, X provides the production-side flexibility that is absent from the AR&DGR model and serves as a diagnostic representation of additional regulation.

**Figure 3.**
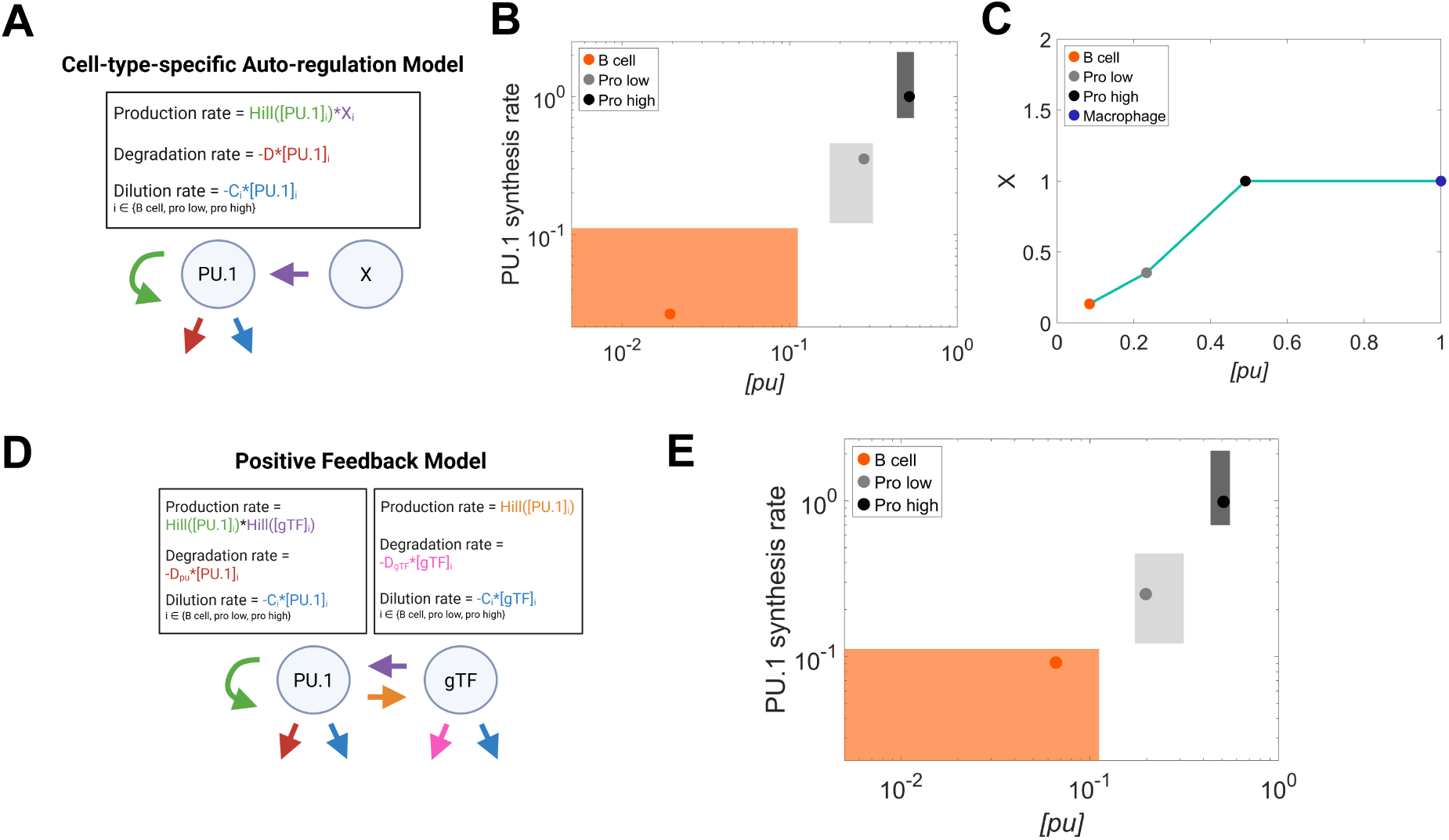
A cell-type-specific input to PU.1 production reproduces experimental PU.1 levels and PU.1 synthesis rates across cell states. **(A)** The architecture of the “cell-type-specific-autoregulation model” is shown. Compared to the “AR&DGR model” in Figure 2A, the model assumes that an additional factor “X” (purple arrow) modulates PU.1 production in a cell-type-specific manner denoted by the subscript “i”. **(B)** Fitted PU.1 abundance and synthesis rates from the model in (A) are compared with the joint measurement ranges. **(C)** Best-fit values of “X” for the three non-reference cell types are plotted against the corresponding PU.1 abundances. **(D)** The architecture of the “positive-feedback model” is shown. The generic TF replaces the independently fitted factor in (A). The generic TF (“gTF”) is produced with a Hill function dependent on PU.1 abundance (orange arrow), degraded at a constant rate (pink arrow), and diluted at a constant rate (blue arrow). **(E)** Fitted PU.1 abundances and synthesis rates from the model in (D) are compared with the joint measurement ranges.

The model reproduced the joint ranges of PU.1 abundance and synthesis rate for all three non-reference cell states (Figure 3B). This result shows that the disagreement identified in the AR&DGR model can be resolved by modulating PU.1 production across cell states. Also, the fitted values of X increased with PU.1 abundance from B cells to pro-low and pro-high cells (Figure 3C). Although this relationship does not uniquely determine the underlying mechanism, it is consistent with a second regulatory factor whose activity increases with PU.1.

We next replaced the independently fitted values of X with an explicit regulatory interaction. In this “positive-feedback model”, PU.1 activates production of a generic transcription factor, gTF, through a Hill function, and gTF in turn increases PU.1 production (Figure 3D). Like PU.1, gTF is removed through natural degradation and dilution during cell division. Its degradation rate was fitted, whereas its dilution rate was constrained by the same cell-cycle measurements used for PU.1. The regulatory functions and their parameters were shared across cell states; the cell states differed through their measured dilution rates.

The positive-feedback model reproduced the measured PU.1 abundance and synthesis-rate ranges for all three non-reference states (Figure 3E). Thus, a second production-side input provides a simple regulatory architecture capable of generating the quantitative differences that PU.1 autoregulation and cell-type-specific dilution could not reproduce alone. C/EBP*α* is a plausible biological candidate for this input. C/EBP*α* binding at the PU.1 promoter and enhancer increases PU.1 expression [13, 17, 22], while PU.1 binding at the *Cebpa* enhancer increases *Cebpa* expression [23]. We therefore asked whether replacing gTF with C/EBP*α* could reproduce the cell-state measurements when the PU.1 production function was also constrained by enhancer-reporter data.

### 2.4 Enhancer-constrained PU.1-C/EBP*α* feedback does not reproduce the progenitor states

To implement C/EBP*α* as the second regulatory factor, we used reporter measurements that quantify how PU.1 enhancer elements and mutations of PU.1 and C/EBP*α* binding sites affect PU.1 expression [13]. These constructs produced distinct reporter activities in macrophages and B cells (Figure 4A; Supplementary Figure S2). Assuming that reporter activity is proportional to PU.1 protein production, we used these measurements to constrain how PU.1 and C/EBP*α* abundance influence the PU.1 production rate.

**Figure 4.**
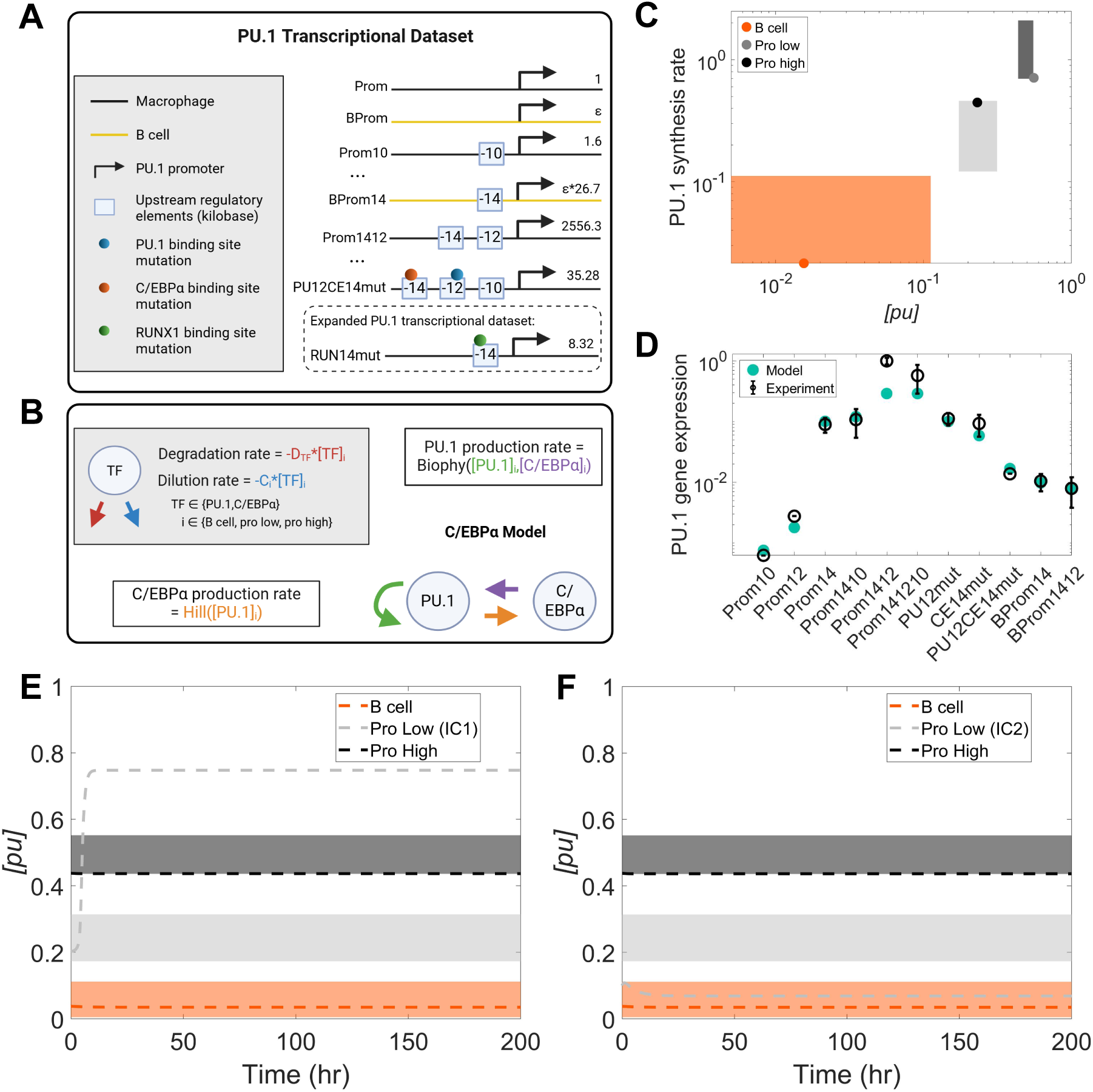
A biophysical model with PU.1-C/EBP*α* positive feedback is insufficient. **(A)** A subset of PU.1 reporter activity measurements used to constrain the PU.1 production rate is shown. A fitted parameter *ε* represents the ratio of reporter activity between the promoter-only construct in B cells (“BProm”) and the promoter-only construct in macrophages (“Prom”). All data points are normalized to the “Pro1412” expression (Methods 5.6). The complete list of constructs and perturbations are described in Table S6 and Supplementary Figure S2. The “expanded PU.1 transcriptional dataset” refers to the same dataset combined with an additional measurement following mutation of RUNX1-binding sites. **(B)** The architecture of the “C/EBP*α* model” is shown. PU.1 production is assumed to be proportional to expression from the wild-type construct, and computed using the statistical-mechanical biophysical function (“Biophy”) constrained by the measurements in (A). Thus, the “PU.1 transcriptional dataset” constrains both PU.1 autoregulation (green arrow) and C/EBP*α*-dependent activation (purple arrow). The “Pro141210” construct was treated as the wild type (Methods 5.6). C/EBP*α* production is a phenomenological function dependent on PU.1 abundance (orange arrow). For each transcription factor (TF) present in the model, its removal rate depends on both degradation (red arrow) and dilution (blue arrow). As before, the dilution rate is cell-type-specific and constrained by measured cell-cycle lengths [5]. **(C)** Fitted PU.1 abundance and production rate are compared with the joint measurement ranges for the “C/EBP*α* model”. **(D)** Fitted reporter activities are compared with the “PU.1 transcriptional dataset” for the “C/EBP*α* model”. The Hill coefficient for C/EBP*α*-dependent activation is 2 in (C) and (D). **(E)-(F)** Cell-state-specific trajectories predicted by the “C/EBP*α* model” are shown for two pro-low initial conditions (ICs), using the best-fit open-loop parameters and a high-cooperativity Hill function. The horizontal colored bands represent the measured PU.1 abundance reported in [5].

We represented PU.1 production using a statistical-mechanical model of enhancer occupancy. The model incorporates two established effects of transcription-factor binding: recruitment of RNA polymerase to the promoter and destabilization of nucleosome formation [24, 25] (Methods 5.5). The reporter measurements constrain this abundance-to-production relationship in macrophages and B cells. We then applied the same relationship to pro-low and pro-high cells, assuming that differences in transcription-factor abundance are sufficient to generate the cell-state-dependent production rates.

The remaining arm of the feedback loop describes activation of C/EBP*α* by PU.1. Because corresponding *Cebpa* reporter measurements were not available, we retained a phenomenological Hill function for this interaction. We initially set the Hill coefficient to two, motivated by the two reported PU.1 binding sites in the *Cebpa* enhancer [23]. The C/EBP*α* degradation rate was constrained by its reported half-life [26], and its dilution rate was determined from the cell-state-specific cell-cycle lengths. We fitted this closed-loop C/EBP*α* model simultaneously to the PU.1 reporter measurements [13] and to the PU.1 abundance and synthesis-rate ranges [5] of the non-reference cell states (Figure 4B). Under these constraints, this “C/EBP*α* model” did not produce a joint fit to the two datasets (Figure 4C, D).

To identify the source of this failure, we opened the feedback loop and fitted the C/EBP*α* abundance of each cell state independently. This open-loop model reproduced both datasets (Supplementary Figure S3), showing that the enhancer-constrained PU.1 production function can accommodate the measurements when appropriate C/EBP*α* levels are supplied. The difficulty therefore arose in generating those C/EBP*α* levels through the PU.1-dependent feedback relationship. In particular, the Hill function with a coefficient of two changed too gradually with PU.1 to reproduce the PU.1-to-C/EBP*α* relationship required by the open-loop fit (Supplementary Section 4). A substantially higher effective cooperativity was needed to recover that relationship (Supplementary Figure S3).

Increasing the cooperativity, however, did not fully resolve the failure of the closed-loop dynamics. We combined the best-fit open-loop parameters with a high-cooperativity PU.1- to-C/EBP*α* relationship and examined the resulting steady states. For the pro-low system, the closed-loop model became bistable, but neither stable state fell within the measured PU.1 abundance range (Figure 4E, F; Supplementary Figure S4). The central limitation was therefore the placement of the attractors rather than simply the ability to generate multiple stable states. Under the biological constraints represented here, PU.1-C/EBP*α* feedback alone does not position the pro-low state correctly. This result motivated us to examine whether an additional enhancer-level input could shift PU.1 production and restore the measured attractor.

### 2.5 RUNX1 input into PU.1-C/EBP*α* feedback leads to predictions consistent with all data constraints

We next searched for an additional regulator acting at the PU.1 enhancer. RUNX1 binds the upstream regulatory region of PU.1, and mutation of its binding sites was reported to reduce PU.1 reporter activity by approximately 90% in macrophages [16]. This large effect suggested that RUNX1 could provide an additional input to PU.1 production. We therefore expanded the biophysical production function to include PU.1, C/EBP*α*, and RUNX1 abundance as inputs (Figure 5A). RUNX1 binding was assumed to influence RNA polymerase recruitment and nucleosome stability through the same general mechanisms used for PU.1 and C/EBP*α*. The production-function parameters were fitted to the original PU.1 reporter measurements together with the RUNX1 binding-site perturbation, collectively referred to as the expanded PU.1 transcriptional dataset (Figure 4A; Methods 5.5).

**Figure 5.**
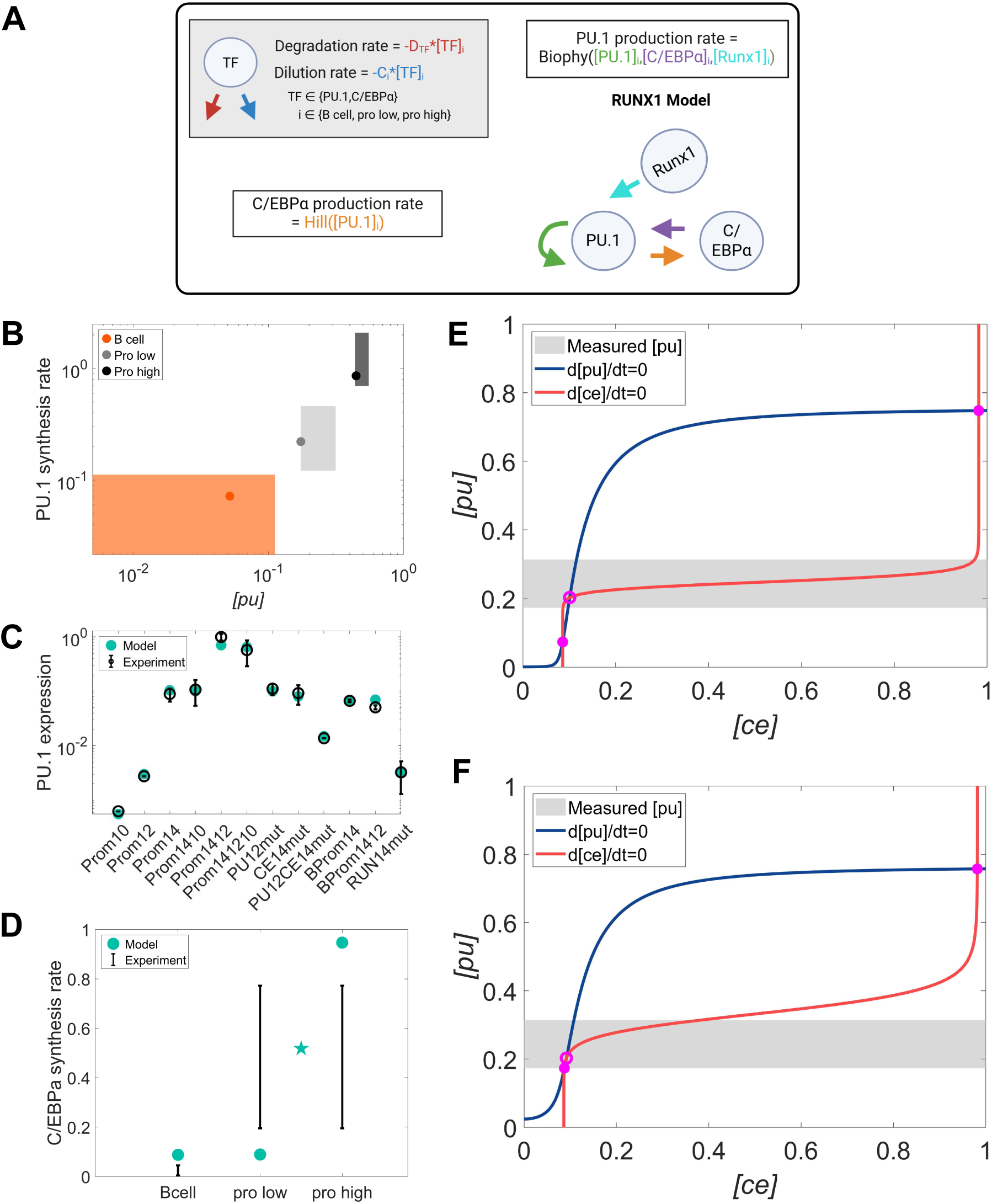
A biophysical model combining PU.1-C/EBP*α* positive feedback with RUNX1 is consistent with measured PU.1 abundance, PU.1 synthesis rates, and PU.1 reporter activity. **(A)** The architecture of the “RUNX1 model” is shown. Relative to the “C/EBP*α* model”, this model includes RUNX1 abundance as an additional input. The PU.1 production function (light-blue, green, and purple arrows) is constrained by the “expanded PU.1 transcriptional dataset” (see Figure 4A). **(B)** Fitted PU.1 levels and synthesis rates from the “RUNX1 model” are compared with the joint measurements. **(C)** Fitted reporter activities from the “RUNX1 model” are compared with the experimental “expanded PU.1 transcriptional dataset”. **(D)** The C/EBP*α* production rates predicted by the “RUNX1 model” are compared with the experimental *Cebpa* RNA-seq measurements [29]. The star marks the mean of the model-predicted C/EBP*α* production rates for pro-low and pro-high. **(E)-(F)** Nullclines for the pro-low system are shown for the high-cooperativity “C/EBP*α* model” and high-cooperativity “RUNX1 model”, under their respective best-fit parameters. The production rate function of the “RUNX1 model” includes an additional enhancer-level input and is also constrained by the PU.1 reporter measurement following RUNX1-site mutation. The intersections of nullclines for all state variables (i.e., PU.1 and C/EBP*α*) represent system steady states. The stability of each steady state is determined using the eigenvalues of the system Jacobian, with an open marker denoting an unstable state and a filled marker denoting a stable state.

The addition of RUNX1 was also consistent with the assumption that cell-state-dependent production arises from differences in transcription-factor abundance. Several studies report distinct RUNX1 levels in B cells, macrophages, and multipotent hematopoietic progenitors [27–29]. We therefore constrained RUNX1 abundance using RNA-seq measurements normalized to the macrophage value, using RNA abundance as a proxy in light of the reported short RUNX1 half-life [29] (Supplementary Section 5.2.3). All other components of the model, including the PU.1-C/EBP*α* feedback and cell-state-specific dilution rates, were retained from the “C/EBP*α* model”.

With a Hill coefficient of two for PU.1-dependent activation of C/EBP*α*, the “RUNX1 model” reproduced the PU.1 abundance and synthesis-rate measurements but not the reporter dataset (Supplementary Figure S5). We therefore allowed the effective cooperativity of this interaction to vary. At a Hill coefficient of ten, the model achieved joint agreement with the cell-state and expanded reporter datasets (Figure 5B, C). This high value should be interpreted as effective cooperativity in the modeled regulatory response rather than as a direct count of PU.1 binding events; possible molecular contributions to this response are considered in the Discussion. Within the model, the combination of RUNX1-sensitive enhancer regulation and a switch-like C/EBP*α* response provides the production-side flexibility needed to satisfy the two sets of constraints.

Nullcline analysis illustrates how RUNX1 improves the placement of the pro-low attractor. In the high-cooperativity C/EBP*α* model, the PU.1 and C/EBP*α* nullclines intersect at a stable state below the measured pro-low PU.1 range (Figure 5E). Incorporating RUNX1 changes the fitted PU.1 production function and shifts the PU.1 nullcline upward at low C/EBP*α* abundance. The resulting intersection falls within the measured pro-low range (Figure 5F). Because the removal function for pro-low is unchanged between the two models, this shift arises from the altered dependence of PU.1 production on enhancer-level regulatory inputs (Supplementary Section 6).

Finally, we compared the model-predicted C/EBP*α* production rates, which were not used during fitting, with *Cebpa* RNA-seq measurements from B cells, macrophages, and multipotent progenitors [29]. The B-cell prediction was slightly above the reported range, and the pro-low and pro-high predictions did not individually match the single progenitor range (Figure 5D). The RNA-seq dataset, however, does not distinguish the two operationally defined progenitor groups. When the two model predictions were averaged, the resulting value (star marker) fell within the measured progenitor uncertainty. The comparison is therefore mixed but broadly consistent with the cell-state trend predicted by the model.

Together, these results show that an additional enhancer-level input represented by RUNX1 can resolve the attractor-placement problem of the “C/EBP*α* model” and produce joint agreement with the PU.1 cell-state and reporter constraints. The analysis supports RUNX1 as a biologically plausible component of the regulatory architecture and identifies enhancer-level control of PU.1 production as a testable mechanism for positioning lineage-associated PU.1 states.

## 3 Discussion

Graded PU.1 abundance is closely associated with hematopoietic stem cell lineage specification. Motivated by an influential dynamical-systems model, we asked whether PU.1 autoregulation and growth-rate feedback can position the reported cell states within their measured ranges rather than merely generate multistability. By evaluating PU.1 abundance, synthesis rate, cell-cycle length, and enhancer-reporter perturbations together, we identified regulatory features needed for quantitative attractor placement within the tested model class. The resulting analysis supports a plausible architecture that combines cell-type-specific dilution, PU.1-C/EBP*α* feedback, and an additional enhancer-level input represented by RUNX1.

The sequence of models separates the contributions of PU.1 removal and production. Measured cell-type-specific dilution corrects the unsupported relationship between PU.1 abundance and cell-cycle length but does not enable PU.1 autoregulation alone to reproduce the progenitor states. PU.1-C/EBP*α* feedback adds production-side flexibility, yet the enhancer-constrained model still misplaces the pro-low attractor. Agreement is recovered when the production function includes an additional enhancer-level input represented by RUNX1. Thus, generating multiple stable states is not sufficient; the architecture must also position attractors correctly under independent cell-state and cis-regulatory constraints.

The model also yields mechanistic predictions for experimental testing. First, bistability in the pro-low system (Figure 5F) indicates the presence of two progenitor subpopulations with different PU.1 abundances but similar growth rates. More broadly, this suggests that additional progenitor cell types may be involved in the HSC differentiation network, beyond the two operationally defined pro-low and pro-high. This prediction is consistent with recent immunophenotyping studies showing that the classical differentiation roadmap can be expanded to include five distinct multipotent progenitor (MPP) groups, known as MPP1- MPP5 [30–32]. A separate study showed that MPP4 comprises two cell populations that differ in cell-cycle lengths, with the slow-and fast-cycling populations each giving rise to distinct lineages [32]. Although the two MPP4 subsets share characteristics with the two progenitor cell states in our model, we lack evidence that pro-low and pro-high correspond to these subsets. Nevertheless, the observed association between progenitor growth rates and lineage output is consistent with our model, in which cell-type-specific growth rates affect lineage specification through their effects on PU.1 abundance.

Within the best-fit model, PU.1-dependent C/EBP*α* production requires high effective cooperativity. Because only two PU.1 binding sites have been reported on the *Cebpa*enhancer [23], the fitted Hill coefficient should not be interpreted as a direct count of PU.1 binding events. Instead, it indicates that additional interactions may contribute to the switch-like activation of *Cebpa* expression. Based on the observed C/EBP*α* binding on its own enhancer, we asked whether C/EBP*α* autoregulation could generate the effective response steepness required by the model (Methods 5.5). An alternative model incorporating this feedback reproduced the data constraints, showing that C/EBP*α* autoregulation is one plausible mechanism (Supplementary Figure S6). Other mechanisms, including the co-condensation of transcription factors and chromatin remodelers, could also contribute to the effective cooperativity [33, 34]. The molecular basis of the effective cooperativity remains to be determined experimentally.

Another mechanistic prediction concerns PU.1 basal gene expression. In our biophysical models, we independently fitted a parameter representing the ratio of promoter-only reporter activity in B cells relative to macrophages (Methods 5.6). We found that this ratio must exceed 1 to reproduce the joint measurements and enhancer perturbation data (Supplementary Figure S7). Within the tested model architecture, the model therefore predicts that promoter-driven basal PU.1 expression is higher in B cells than in macrophages. Given that the B cell production rate is lower than that in macrophages, this suggests that cell-type differences in protein production rate cannot be inferred from basal expression alone, but is shaped by additional transcriptional regulation. Further perturbation studies are needed to test this prediction.

Several limitations should be considered when interpreting the broader relevance of our findings. Although all measurements used to define the biological constraints were obtained from mouse studies, we did not explicitly account for cross-study differences. We combined distinct measurement modalities, including GFP-based measurements of PU.1 abundance and synthesis, luciferase reporter activity, and RNA abundance. These quantities constrain related aspects of the regulatory system but are not directly equivalent. In addition, RNA abundance was used as a proxy for protein production, and DNA occupancies inferred from reporter assays may not reflect endogenous chromatin regulation at the PU.1 locus during differentiation. Matched measurements collected from the same defined cell populations would reduce these uncertainties.

The modeling framework introduces additional limitations. Stable fixed points provide a useful representation of distinct cell populations but do not capture noise or transient states along the differentiation trajectory. Incorporating RUNX1 also increases the model’s flexibility, at the cost of reduced parameter identifiability. Finally, the aggregate progenitor RNA-seq data used for external evaluation cannot independently test the pro-low and pro-high predictions for C/EBP*α* production. Our conclusions are therefore specific to the integration of these datasets under the chosen assumptions and do not imply a unique biological requirement for C/EBP*α* and RUNX1.

Overall, our study combines dynamical systems theory and statistical mechanics to examine how transcription-factor abundance and enhancer regulation shape lineage-associated PU.1 states. The framework constrains system attractors using steady-state measurements of lineage-specific protein abundance and production. By requiring the same model to reproduce enhancer-perturbation data, the analysis generates testable biophysical hypotheses for enhancer-mediated regulation of PU.1 expression. More broadly, this framework could be applied to other differentiation systems in which master transcription factors help shape lineage-associated states, such as myogenic differentiation controlled by MyoD1 [35] and embryonic stem cell fate commitment by Oct-3/4 [36].

## 4 Methods

### 4.1 Normalization of HSC differentiation dataset

Measurements of PU.1-GFP level, PU.1 synthesis rate, and cell-cycle length in B cells, pro-low cells, pro-high cells, and macrophages were extracted using PlotDigitizer [37] from Figure 1H of [5]. We hereafter refer to these data as “HSC differentiation dataset”. All models were fitted to the joint PU.1-GFP and synthesis-rate measurements of this dataset, whereas the cell-cycle lengths were used to precompute the protein removal rates. For PU.1-GFP intensity and PU.1 production rate, we extracted the reported median and central 50% of each distribution; for cell-cycle length, we extracted the reported mean and 95% confidence interval.

We then normalized each data type by the corresponding macrophage measurement. We assumed that the experimentally reported “protein synthesis rate” corresponds to the model-predicted “protein production rate”. These terms are used interchangeably throughout the manuscript.

The measurement uncertainties are asymmetric, so the standard error-propagation formula derived using a Taylor expansion is not applicable. In Supplementary Section 1, we describe an alternative workflow for propagating uncertainty during normalization. For each measurement type, the resulting uncertainty interval is shown as a colored patch in Figures 1C and 1D.

To incorporate earlier evidence on PU.1-guided HSC differentiation, we also considered measurements from a previous study [4]. That study helped establish the lineage-specific effects of PU.1. Specifically, Figure 2A of [4] shows the PU.1-GFP distributions for macrophages and pro-B cells derived from PU.1-homozygous-knockout mice and rescued by PU.1 transduction.

We took the median of each cell-type distribution [4] and computed the B cell-over- macrophage PU.1-GFP ratio. This ratio is approximately 1/200, about 10-fold smaller than the propagated lower uncertainty bound obtained using the preceding procedure [5]. We used this smaller value as the normalized lower bound of the B-cell PU.1-GFP interval.

### 4.2 Dimensionless analysis of the “AR&GF model”

To examine the “AR&GF model”, we first replicated it based on Equations 23, 25, and 26 in the section titled “Phase Plane Analysis of the Different Feedback Architectures” in the Supplementary Materials of [5]. The ordinary differential equation (ODE) for the “AR&GF model” [5] is:

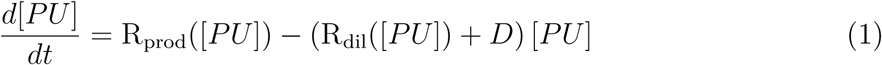

Here, the notations R_prod_(*x*) and R_dil_(*x*) refer to the production rate and the dilution rate constant, given input *x*:

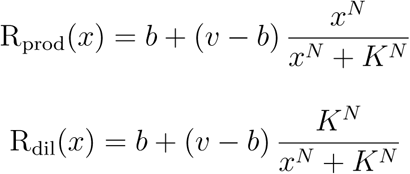

where *v*, *b*, *N*, and *K* are the maximum rate (constant), basal rate (constant), Hill coefficient, and half-saturation constant, respectively.

According to Eqn. (1), PU.1 concentration, [*PU*], increases at a rate modeled by an increasing Hill function of [*PU*]. PU.1 is diluted at a rate that also depends on [*PU*] and is degraded with rate constant *D*.

Using Eqn. (1) and parameters from Table S6 of [5], we reproduced the model shown in the lower panel of Figure 4 in [5] (not shown). The “dilution rate” shown there plots (R_dil_([*PU*]) + *D*)[*PU*] as a function of [*PU*].

To avoid confusion, we use “removal rate” for this model component (Figure 1), defined as the sum of the protein dilution rate (due to cell division) and the protein degradation rate (due to natural decay).

Next, to compare model predictions with normalized data, we nondimensionalized Eqn. (1) using its highest stable steady state (the macrophage state) (see Supplementary Section 2.1). After nondimensionalizing, the “AR&GF model” is as follows:

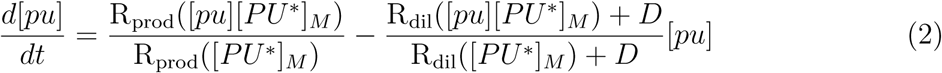

where [*PU ^∗^*]*_M_* is the highest steady-state solution, or the macrophage state, of Eqn. (1), and [*pu*] = [*PU*]*/*[*PU ^∗^*]*_M_* is the dimensionless concentration. Because [*PU ^∗^*]*_M_* is not directly reported in [5], we identified it by finding the largest [*PU*] at which production and removal rates are equal.

Strictly, nondimensionalizing Eqn. (1) also introduces a constant prefactor, 1*/* (R_dil_([*PU ^∗^*]*_M_*) + *D*), on the left-hand side of Eqn. (2) (see Supplementary Section 2.1). As such, time in Eqn. (2) is not dimensionless. We omitted this factor because it is a positive constant that does not affect the locations of the steady states or their stability, which is the main focus of our work.

Because the highest stable steady state (the macrophage state) with the largest production rate was used for normalization, both [*pu*] and the dimensionless production rate, R_prod_([*pu*][*PU ^∗^*]*_M_*)*/*R_prod_([*PU ^∗^*]*_M_*), range from 0 to 1. Therefore, the point (1, 1) in all main-text figures (star marker) represents the macrophage reference state and is not an independent prediction.

### 4.3 General modeling framework with cell-type-specific growth rates

We used the same modeling principles as Eqn. (1) to describe the rate of change in PU.1. Because we assigned a cell-type-specific dilution rate from the measured cell-cycle length, we modeled the rate of change in PU.1 for each cell type with a separate ODE. Thus, the models can be expressed generally as:

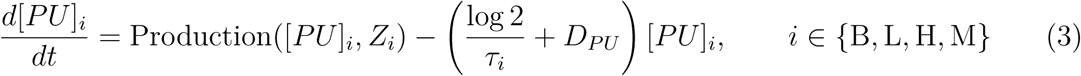

where B, L, H, M denote B cell, pro-low, pro-high, and macrophage.

For cell type *i*, PU.1 is produced at rate Production([*P U*]*_i_, Z_i_*). Because all models include PU.1 autoregulation, this function depends on the PU.1 abundance of that cell type. *Z_i_*is a vector accommodating additional inputs (in this case, abundance of other transcription factors). Within the same model, parameters of Production([*P U*]*_i_, Z_i_*) are shared across cell types unless explicitly subscripted by *i*. The removal-rate constant contains the dilution term log(2)*/τ_i_*, where *τ_i_* is the cell-type-specific cell-cycle length, and the degradation constant *D_PU_* . The factor log(2) follows from the assumption of exponential cell growth [38].

We then nondimensionalized Eqn. (3) using the macrophage steady state, defined by *d*[*PU*]*_M_ /dt* = 0, together with the steady-state conditions of any additional transcription factors in *Z_M_*, which gives:

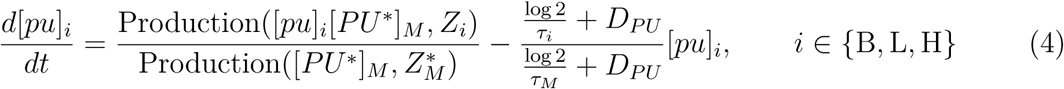

Here, [*pu*]*_i_* = [*PU*]*_i_/*[*PU ^∗^*]*_M_* is the cell-type-specific dimensionless PU.1 concentration. *Z_M_\** contains the abundances of the non-PU.1 transcription factors at the macrophage steady state. *τ_M_* is the macrophage cell-cycle length. The PU.1 degradation constant is *D_PU_* = 1*/*70 h*^−^*^1^, based on the average PU.1 decay rate across two experiments in Figure S6 of [5]. We here again omitted the constant prefactor on the left-hand side of Eqn. (4).

For all subsequent models, we precomputed the dimensionless removal-rate ratio [(log 2)*/τ_i_* + *D_PU_*]*/*[(log 2)*/τ_M_* + *D_PU_*]. Because *τ_i_*is uncertain, this ratio is also uncertain. Using an error-propagation procedure analogous to that in Methods 5.1, we obtained a distribution of dimensionless removal-rate constants. The mean of the error-propagated ratio was used in Eqn. (4) except for pro-high cells, for which the 25th percentile was used. Because this ratio depends on the inverse of the cell-cycle length, the 25th percentile of the ratio was approximately computed with the upper (75th) percentile of the pro-high cell-cycle lengths. Detailed steps are provided in Supplementary Section 2.2; the rationale and corresponding analysis are provided in Supplementary Section 2.3 and Supplementary Figure S8.

### 4.4 Formulation of phenomenological models

Because the PU.1 removal-rate constant was precomputed across models, the following formulations specify only Production([*P U*]*_i_, Z_i_*). For models without additional transcription-factor inputs (Eqns. (6) and (7)), *Z_i_*is empty and omitted. For additional state variables, which here are other transcription factors, we provide the complete ODE. Although all models were implemented in the dimensionless form of Eqn. (4), for simplicity we show the production terms in the dimensional form of Eqn. (3). Supplementary Section 2.4 provides further details, and Table S1 compares all phenomenological models.

We first define a *dimensionless* Hill function to be referred to throughout all phe- nomenological formulations:

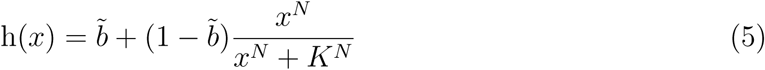

The parameters *b*, *N*, and *K* are specific to each Hill function: every occurrence of h(*·*) in the models below has its own *b͂*, *N*, and *K*, even when two occurrences share the same input. Here, *b͂* is the dimensionless basal level, distinct from the dimensional basal rate *b* in R_prod_(*x*).

Given any input *x*, the output of h(*x*) does not exceed 1. In the following models, h(*x*) may be multiplied by a scaling factor to be consistent with the unit of the corresponding equation.

#### 4.4.1 Autoregulation and distinct growth rate (AR&DGR) model

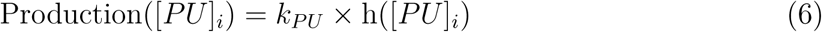

Here, *k_PU_* is the dimensional production rate.

#### 4.4.2 Cell-type-specific-autoregulation model

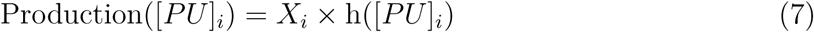

Here, *X_i_*is a cell-type-specific dimensional production rate.

#### 4.4.3 Positive feedback model

This model consists of two ODEs. The first, describing the change in PU.1 abundance, has the form

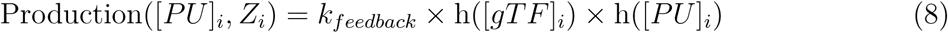

Here, *Z_i_* = [*gTF*]*_i_*, and *k_feedback_* is the dimensional production rate. Now the production rate of PU.1 is additionally activated by the generic TF (“gTF”) abundance in a Hill manner, and the cell-type-specificity of production rate comes from h([*gTF*]*_i_*).

The second ODE describes the change in gTF abundance, which is activated by PU.1:

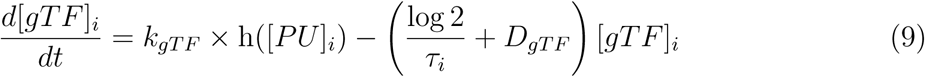

Here, cell-type-specific [*gTF*]*_i_* is produced at a rate modeled by an increasing Hill function of PU.1 and removed by both dilution and degradation. *k_gT F_* is the dimensional production rate. *τ_i_* is the same cell-type-specific cell-cycle length used to constrain PU.1 removal, and *D_gT F_* is the generic-TF degradation-rate constant.

### 4.5 Formulation of biophysical models

As in the phenomenological models, the governing equation for the changes in protein concentration depends on production and removal rates. Here, however, the PU.1 production function is additionally constrained by PU.1 reporter activity. Detailed biophysical model formulations are provided in Supplementary Sections 2.5.1–2.5.2. A comparison between all biophysical models is shown in Table S2.

#### 4.5.1 C/EBP*α* model

In the “C/EBP*α* model”, the PU.1 production rate function is

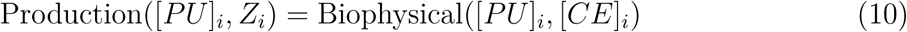

with *Z_i_* = [*CE*]*_i_*.

We assumed that PU.1 expression is regulated by PU.1 and C/EBP*α* binding at three enhancer elements located 14, 12, and 10 kilobase pairs upstream of the PU.1 promoter [13]. PU.1 production was modeled as directly proportional to gene expression through the function Biophysical([*PU*]*_i_,* [*CE*]*_i_*). Its detailed formulation is provided in Supplementary Sections 2.5.3–2.5.4, and Tables S3–S5 list the possible occupancies of the three enhancer elements (by PU.1 and C/EBP*α*) and promoter (by RNAP).

Briefly, we modeled two transcriptional regulatory mechanisms that can affect PU.1 gene expression: (i) TF binding can directly recruit or repel transcriptional machinery; (ii) TF binding can destabilize nucleosomes at other TF-binding sites and at the promoter. In addition, the biophysical models account for protein–protein interactions between co-bound TFs. Evidence supporting these mechanisms and interactions mediated by PU.1 and C/EBP*α* is summarized in Table S8.

The ODE describing the change in C/EBP*α* level is

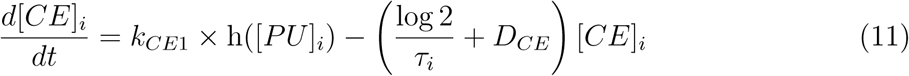

Here, C/EBP*α* is produced in a Hill manner dependent on PU.1 abundance with *k_CE_*_1_ being the dimensional production rate. The Hill coefficient *N* in h([*PU*]*_i_*) was initially constrained by the number of PU.1 binding sites (two) on the *Cebpa* enhancer [23]. *D_CE_* is the natural degradation-rate constant of C/EBP*α* and was set to 0.4159 h*^−^*^1^. This value was calculated from a reported half-life of approximately 100 minutes [26]. In the dimensionless model, the removal-rate ratio for C/EBP*α* was obtained using the same error-propagation procedure as the corresponding ratio for PU.1, as described in Methods 5.3.

The main text also considers an “open-loop C/EBP*α* model”. This model contains Eqn. (10), but rather than computing C/EBP*α* with an ODE, we treated [*CE*]*_i_* for *i ∈ {*B, L, H*}* as unknown parameters (Table S7).

Our model focuses on PU.1 transcriptional regulation by the upstream enhancer. PU.1 expression is also regulated by other regulatory elements. In particular, transcription factors Oct-1, Oct-2, Sp1, and PU.1 bind to the murine PU.1 promoter and contribute to cell-type-specific expression between macrophages and B cells [14, 39]. However, after mutation of these TF-binding sites, the fold change in PU.1 expression (less than 10^1^) is much smaller than the change resulting from mutations of enhancer TF-binding sites (at least 10^2^) [13, 15, 40]. More importantly, gene constructs carrying only the PU.1 promoter in transgenic models cannot induce PU.1 expression [15]. We therefore focused primarily on the effect of the upstream enhancer on PU.1 expression.

#### 4.5.2 RUNX1 model

The RUNX1 model follows the same principles as the “C/EBP*α* model”. The equation describing the rate of change in C/EBP*α* is the same as Eqn. (11). The biophysical PU.1 production rate additionally depends on RUNX1 abundance. This extension was motivated by the observation that PU.1 expression is affected by perturbation of RUNX1 binding sites [16]:

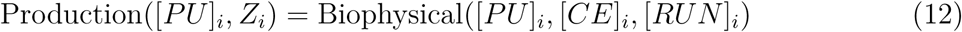

with *Z_i_* = ([*CE*]*_i_,* [*RUN*]*_i_*).

The model incorporates the effects of PU.1, C/EBP*α*, and RUNX1 on RNA polymerase occupancy. In this model, RUNX1 was likewise assumed to interact with RNA polymerase and destabilize nucleosome formation. Tables S3–S5 list all possible occupancies on the three enhancer elements by PU.1, C/EBP*α*, and RUNX1, and promoter by RNAP. Evidence supporting these RUNX1-mediated mechanisms is summarized in Table S8.

One of the three reported RUNX1 binding sites [16] overlaps a C/EBP*α* binding site (C1 in [22]) in the proximal homology region of the PU.1 enhancer. The decrease in PU.1 expression after RUNX1-site mutation could therefore partly reflect loss of C/EBP*α* binding. However, mutation of C1 reduced PU.1 expression by approximately 20% (from 100% to 77%; Figure 3C in [22]), whereas mutation of all RUNX1 sites reduced expression by approximately 90% (from 870% to 86%; Figure 1D in [16]). Thus, the nonoverlapping RUNX1 sites make a substantial contribution, supporting a RUNX1 effect that is distinct from C/EBP*α* regulation.

#### 4.5.3 RUNX1 model with C/EBP*α* autoregulation

The best-fit “RUNX1 model” required high cooperativity for PU.1-dependent activation of C/EBP*α* (*N* = 10 in Eqn. (11)). Because this value exceeds the two reported PU.1 binding sites on the *Cebpa* enhancer [23], we asked whether C/EBP*α* autoregulation [23, 41] could generate the required effective response. We therefore developed a third biophysical model with low-cooperativity PU.1 activation and Hill-type C/EBP*α* autoregulation:

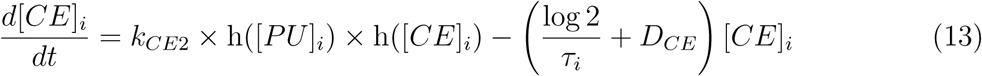

where *k_CE_*_2_ is the dimensional production rate. Details of the model constraints can be found in Table S8.

### 4.6 Fitting procedures

The dimensionless formulation in Eqn. (4) contains the dimensional macrophage steady state [*PU ^∗^*]*_M_* . Rather than fitting it as an independent parameter, we reparameterized each model so that it was absorbed into existing parameters. Supplementary Section 3 describes this normalization, and Supplementary Section 4 shows how the normalized open-loop C/EBP*α* model informs the cooperativity needed in the closed-loop model.

All models were implemented in MATLAB. For models without positive feedback, each model comprised three ODEs describing the rate of change in PU.1 level. Each ODE corresponded to a cell type and differed only in the removal rate. The analysis included the three non-reference cell types. Similarly, for models with feedback, we used three ODE systems, with each system containing two ODEs. The first ODE described the rate of change in PU.1, and the second the rate of change in the generic TF or C/EBP*α*.

We fitted all models to the joint PU.1-GFP and synthesis-rate measurements for non-reference cell types in the “HSC differentiation dataset”, using an objective function that penalized model-predicted steady-state PU.1 concentrations outside the experimental PU.1-GFP ranges. The function also penalized predicted PU.1 production rates outside the experimental synthesis-rate ranges. The complete procedure for fitting all models to the joint PU.1-GFP and synthesis-rate measurements is provided in Supplementary Section 5.1.

For biophysical models, the PU.1 production rate was additionally constrained by PU.1 reporter activity data. Specifically, the “C/EBP*α* model” was fitted to PU.1 expression measurements upon PU.1 and C/EBP*α* binding site perturbations (“PU.1 transcriptional dataset” [13]). The “RUNX1 model” was fitted to PU.1 expression measurements following perturbations of PU.1, C/EBP*α*, and RUNX1 binding sites (“expanded PU.1 transcriptional dataset” [13] and [16]). A full description of the gene constructs in both datasets is provided in Table S6. The detailed procedure for processing these gene expression datasets is provided in Supplementary Section 5.2.1. The procedure for fitting each model to its respective gene expression dataset is provided in Supplementary Section 5.2.2.

Notably, in the formulation of biophysical models, the measured PU.1 reporter activity constrains the changes in PU.1 production rate within the same cell type. However, these measurements do not determine the relationship between production rates across cell states. We therefore introduced a rescaling parameter (*ε*) to represent the ratio of gene expression between the promoter-only construct in B cells (“BProm”) and the promoter-only construct in macrophages (“Prom”) (Figure 4A). Treating the macrophage “Prom” as the reference measurement, all B-cell measurements were expressed in terms of this factor. *ε* was fitted independently for all biophysical models.

We also imposed quantitative constraints on RUNX1 levels in Eqn. (12) using the “Haemopedia-Mouse-RNASeq” dataset [29], which reports RNA-seq measurements along the hematopoietic stem cell differentiation roadmap, organized by cell lineages and cell types. The detailed formulation of such a constraint is provided in Supplementary Section 5.2.3. The same dataset was used to evaluate the model predictions of C/EBP*α* production rate. We did not constrain C/EBP*α* production rate using RNA-seq data during fitting because no model reproduced C/EBP*α* RNA-seq data. We therefore used the C/EBP*α* production rate data only for external evaluation.

Finally, we assumed that the PU.1 steady states across all cell types arise from the wild-type PU.1 gene. Given the available data, we treated the gene construct corresponding to the “Prom141210” data point as the wild type (see Table S6). As such, the biophysical production rate functions in Eqn. (10) and Eqn. (12) assume DNA occupancies consistent with the gene composition of “Prom141210” and compute gene expression using cell-type-specific TF concentrations (Supplementary Section 5.2.2).

All models were fitted using MATLAB particle-swarm optimization [42]. Table S7 summarizes the all known and unknown parameters. Table S8 provides the qualitative and quantitative constraints for all biophysical models.

## Data availability statement

The data and software used for the phenomenological and biophysical models, together with the scripts used to generate the figures, are available at REPOSITORY_URL_OR_ZENODO_DOI.

## Supplemental data

Supplemental data are available online.

## Author contributions

O.I. and Y.Z. conceived the study. Y.Z. analyzed the experimental data and developed and implemented the models. O.I. and Y.Z. wrote and revised the manuscript.

## Acknowledgments

This work was supported by the National Science Foundation under Grant MCB-2204402 and the Welch Foundation under Grant C-1995 to Oleg Igoshin.

