## Supplementary Information for "Integrating cell-state and enhancer constraints discriminates models of PU.1 regulation during hematopoietic differentiation"

**6 Understanding model nullclines** **28**

**7 Supplementary Figures** **29**

### 1 Normalization of the HSC differentiation dataset

Our work is motivated by the PU.1-GFP, PU.1 synthesis rates, and cell-cycle length measurements provided by [1] (hereafter, "HSC differentiation dataset"). All models are fitted to the joint PU.1-GFP and synthesis-rate measurements of this dataset, whereas the cell-cycle lengths are used to precompute the protein removal rates (Section 2.2). In the main text, this dataset is normalized with respect to the macrophage measurement. In this section, we demonstrate the normalization procedures.

Given the asymmetrical uncertainty in the given measurement [1] and the lack of original data, we decide to create probability distributions with the statistics that match the given measurement, and conduct error propagation using sampling.

For all methods below, let $X^i_{mt}$ be a specific measurement with uncertainty, where $i$ (cell type) can be B cells, pro-low cells, pro-high cells, or macrophages, i.e., $i \in \{\mathrm{B}, \mathrm{L}, \mathrm{H}, \mathrm{M}\}$. Additionally, $mt$ (measurement type) can be PU.1-GFP abundance, PU.1 synthesis rate, or cell-cycle length, i.e., $mt \in \{GFP, syn, cycle\}$.

#### 1.1 Finding probability distributions with matching statistics

To find a probability distribution that produces the same statistics as the measurement, we use $X^B_{GFP}$ as an example, with known median and the central 50% of the measurement. Let the measured median, the 25th percentile, and the 75th percentile be $X^B_{GFP}|_{median}$, $X^B_{GFP}|_{q25}$, $X^B_{GFP}|_{q75}$.

Due to an observed skew in measurements, we assume that $X^B_{GFP}$ follows a log-normal distribution. Let this probability distribution be $P^B_{GFP}$, with mean, standard deviation, the 25th percentile, and the 75th percentile being $\mu$, $\sigma$, $Q25$, $Q75$.

By definition, the probability distribution $P^B_{GFP}$, when transformed logarithmically, becomes a normal distribution. Let the log-transformed distribution be $P^B_{GFP,0}$, with statistics $\mu_0$, $\sigma_0$, $Q25_0$, $Q75_0$. The statistics are related to each other using Z score:

$$Q25_0 = \mu_0 - 0.67449 \times \sigma_0 \tag{S1}$$

and

$$Q75_0 = \mu_0 + 0.67449 \times \sigma_0 \tag{S2}$$

Following the transformation $P^B_{GFP,0} = \log(P^B_{GFP})$, we have that:

$$Q25_0 = \log(Q25) \tag{S3}$$

$$Q75_0 = \log(Q75) \tag{S4}$$

Meanwhile, assuming that the measurements also follow a log-normal distribution, then:

$$Q25 = X^B_{GFP}|_{q25} \tag{S5}$$

and

$$Q75 = X^B_{GFP}|_{q75} \tag{S6}$$

From Eqns. (S3)–(S6), we obtain that

$$Q25_0 = \log(X^B_{GFP}|_{q25}) \tag{S7}$$

and

$$Q75_0 = \log(X^B_{GFP}|_{q75}) \tag{S8}$$

Using Eqns. (S1) and (S2), we can solve for the parameters $\sigma_0$ and $\mu_0$ defining the normal distribution using reported measurements as follows:

$$\sigma_0 = \frac{Q75_0 - Q25_0}{2 \times 0.67449} = \frac{\log(X^B_{GFP}|_{q75}) - \log(X^B_{GFP}|_{q25})}{2 \times 0.67449} \tag{S9}$$

$$\mu_0 = \frac{Q75_0 + Q25_0}{2} = \frac{\log(X^B_{GFP}|_{q75}) + \log(X^B_{GFP}|_{q25})}{2} \tag{S10}$$

The distribution describing $X^B_{GFP}$ is then

$$X^B_{GFP} \sim P^B_{GFP} = e^{P^B_{GFP,0}(\sigma_0,\mu_0)} \tag{S11}$$

For measurements reported with the 95% confidence interval, the same method applies except that the Z-score is 1.96 in Eqns. (S1) and (S2). Note that we assume that the bounds of the confidence interval correspond to the 2.5th and 97.5th percentiles of the underlying distribution of the measurement.

#### 1.2 Error propagation using sampling

Now that the probability distribution is obtained for each measurement $X^i_{mt}$, we use the following workflow to propagate uncertainty. For each $X^i_{mt}$:

1. Randomly sample from the corresponding distribution $P^i_{mt}$ to obtain a realization of the measurement, called $simX^i_{mt}$. This can be done using the MATLAB command lognrnd(mu,sigma), where "mu" and "sigma" correspond to $\mu_0$ and $\sigma_0$ in Section 1.1

2. Compute the following ratio, if $simX$ have the same $mt$ between the numerator and denominator, for all possible $i$

$$R^i_{mt} = \frac{simX^i_{mt}}{simX^M_{mt}} \tag{S12}$$

   Each $R^i_{mt}$ is a macrophage-normalized realization of cell type $i$ and measurement type $mt$. For example, $R^B_{GFP} = \frac{simX^B_{GFP}}{simX^M_{GFP}}$ is the macrophage-normalized PU.1-GFP in B cells

3. Repeat Steps 1 and 2 many times

Doing so yields sufficient samples to construct a sample space of $R^i_{mt}$ for all non-macrophage $i$ and $mt$, including $R^B_{GFP}$, $R^L_{GFP}$, $R^H_{GFP}$, ..., $R^H_{syn}$, ..., and $R^H_{cycle}$. For each non-macrophage sample space, we then compute the relevant statistics (median and central 50% of the data, or mean and 95% confidence interval) for $R^i_{mt}$.

There is one exception to this procedure. For the macrophage-normalized B-cell PU.1-GFP abundance ($R^B_{GFP}$), we do not use the error-propagated lower bound. Instead, based on an independent measurement of PU.1-GFP in B cells and macrophages, we set the normalized lower bound to approximately 1/200, about 10-fold smaller than the error-propagated lower bound. The rationale and data source are described in Methods 5.1 of the main text.

### 2 Model development

To be consistent with the normalized experimental data [1], this section includes the steps to nondimensionalize the model developed by the same study (the "AR&GF model" [1]). The detailed explanation of the "AR&GF model" can be found in Methods 5.2 of the main text. In the subsequent sections, we focus on showing model formulations of phenomenological and biophysical models.

#### 2.1 Nondimensionalization of the AR&GF model

Given that PU.1 abundance is modeled with the following ODE in the "AR&GF model":

$$\frac{d[PU]}{dt} = \mathrm{R_{prod}}([PU]) - (\mathrm{R_{dil}}([PU]) + D)\,[PU] \tag{S13}$$

one can convert this ODE to a dimensionless form using the following method [2].

Let the macrophage steady state of $\frac{d[PU]}{dt}$ be $[PU^*]_M$. The steady state solution must satisfy

$$\frac{d[PU]}{dt} = 0 \tag{S14}$$

or

$$\mathrm{R_{prod}}([PU]) = (\mathrm{R_{dil}}([PU]) + D)\,[PU] \tag{S15}$$

Consequently,

$$[PU^*]_M = \frac{\mathrm{R_{prod}}([PU^*]_M)}{\mathrm{R_{dil}}([PU^*]_M) + D} \tag{S16}$$

Define the dimensionless PU.1 abundance as

$$[pu] = [PU]/[PU^*]_M \tag{S17}$$

and rewrite Eqn. (S13) using Eqn. (S17):

$$[PU^*]_M \frac{d[pu]}{dt} = \mathrm{R_{prod}}([pu][PU^*]_M) - (\mathrm{R_{dil}}([pu][PU^*]_M) + D)\,[pu][PU^*]_M \tag{S18}$$

Notably, hereafter, for all transcription factors mentioned in the model, uppercase letters inside of "[]" will denote dimensional abundance, whereas lowercase letters denote dimensionless abundance.

Now rearrange Eqn. (S18) such that the ODE is expressed in terms of the dimensionless PU.1 abundance $[pu]$:

$$\frac{d[pu]}{dt} = \frac{\mathrm{R_{prod}}([pu][PU^*]_M)}{[PU^*]_M} - (\mathrm{R_{dil}}([pu][PU^*]_M) + D)\,[pu] \tag{S19}$$

Given Eqn. (S16), this becomes:

$$\frac{d[pu]}{dt} = \frac{\mathrm{R_{prod}}([pu][PU^*]_M)}{\frac{\mathrm{R_{prod}}([PU^*]_M)}{\mathrm{R_{dil}}([PU^*]_M)+D}} - (\mathrm{R_{dil}}([pu][PU^*]_M) + D)\,[pu] \tag{S20}$$

which can be rearranged as:

$$\frac{1}{\mathrm{R_{dil}}([PU^*]_M) + D}\frac{d[pu]}{dt} = \frac{\mathrm{R_{prod}}([pu][PU^*]_M)}{\mathrm{R_{prod}}([PU^*]_M)} - \frac{\mathrm{R_{dil}}([pu][PU^*]_M) + D}{\mathrm{R_{dil}}([PU^*]_M) + D}[pu] \tag{S21}$$

Although Eqn. (S21) is not fully dimensionless due to the unit of time, given that we are not concerned with the time taken to reach steady state, it is sufficient for our analysis.

#### 2.2 Development of models with cell-type-specific protein removal rate

In general, all models developed in our work follow the same principles as Eqn. (S13) to model the change in PU.1 abundance:

$$\frac{d[PU]}{dt} = \mathrm{Production}([PU], Z) - (\mathrm{Dilution}([PU]) + D_{PU})\,[PU] \tag{S22}$$

where Production and Dilution are functions describing the production rate and dilution rate constant (due to cell division), respectively, and $Z$ is a vector accommodating additional inputs to the production rate (in this case, abundances of other transcription factors). We define the removal rate constant, $\mathrm{Dilution}([PU]) + D_{PU}$, as the sum of the dilution rate constant $\mathrm{Dilution}([PU])$ and the degradation rate constant $D_{PU}$.

Given that we do not observe a monotonic, continuous function describing the relationship between $[PU]$ and the cell-cycle length, we decide to assume that each cell type has a constant, unique cell-cycle length. As such, $\mathrm{Dilution}([PU])$, or the dilution rate constant of intracellular protein, can be estimated from the cell-cycle length as follows:

$$\mathrm{Dilution}([PU]) = \frac{\log 2}{\tau_i} \tag{S23}$$

where $\tau_i$ is the cell-cycle length in cell type $i$ from the “HSC differentiation dataset”. This assumes protein abundances are diluted at the same rate as cells grow exponentially.

Due to the cell-type-specific dilution rates, we work with cell-type-specific ODEs:

$$\frac{d[PU]_i}{dt} = \mathrm{Production}([PU]_i, Z_i) - \left(\frac{\log 2}{\tau_i} + D_{PU}\right)[PU]_i \tag{S24}$$

with $i \in \{\mathrm{B}, \mathrm{L}, \mathrm{H}, \mathrm{M}\}$.

Nondimensionalizing this model using $[pu]_i = [PU]_i/[PU^*]_M$ and the macrophage steady state, defined by $d[PU]_M/dt = 0$ together with the steady-state conditions of any additional transcription factors in $Z_M$, yields the dimensionless form:

$$\frac{d[pu]_i}{dt} = \frac{\text{Production}([pu]_i[PU^*]_M, Z_i)}{\text{Production}([PU^*]_M, Z^*_M)} - \frac{\frac{\log 2}{\tau_i} + D_{PU}}{\frac{\log 2}{\tau_M} + D_{PU}}[pu]_i \tag{S25}$$

with $i \in \{\text{B}, \text{L}, \text{H}\}$, where $Z^*_M$ contains the abundances of the non-PU.1 transcription factors at the macrophage steady state. The steady state of $\frac{d[pu]_M}{dt}$ is 1 and need not be solved; it is thus not considered during model development.

For all model formulations below, we precompute the dimensionless removal rate constant, $\frac{\frac{\log 2}{\tau_i}+D_{PU}}{\frac{\log 2}{\tau_M}+D_{PU}}$, for each cell type. Let this constant be $R^i_{RemRate}$, and it can be obtained using the same normalization and error-propagation procedure in Section 1.2, except that the ratio in Step 2 is now defined as

$$R^i_{RemRate} = \frac{\frac{\log 2}{simX^i_{cycle}} + D_{PU}}{\frac{\log 2}{simX^M_{cycle}} + D_{PU}} \tag{S26}$$

where we use $D_{PU} = 1/70\,\text{h}^{-1}$, based on the average PU.1 decay rate across two experiments in Figure S6 of [1]. The **mean** of each distribution of $R^i_{RemRate}$ is fed into the model.

#### 2.3 Justifications for using a lower removal rate constant of pro-high cells

Notably, we used the 25th percentile, rather than the mean, of the $R^H_{RemRate}$ sample space as the dimensionless removal rate constant for $\frac{d[pu]_H}{dt}$. This section explains why we made this exception.

The dimensionless removal rate constant is assumed to be the mean of $R^i_{RemRate}$, referred to as $meR^i_{RemRate}$ for convenience. We can compute a range of dimensionless removal rates ($meR^i_{RemRate} \times [pu]_i$) at steady state across various PU.1 abundances ($[pu]_i$). Since we are interested in developing models predicting PU.1 steady states that lie within the uncertainty of measured PU.1-GFP abundances, we examined the removal rate evaluated at the 25th and 75th percentiles of the PU.1-GFP measurement, or $R^i_{GFP}$ sample space, obtained from Section 1.2.

For pro-high cells ($i = H$), in the regime where the model produces steady state consistent with data, we found that the minimum dimensionless removal rate is greater than 1 (Figure S8, red triangle marker). At steady state, production rate is equal to the removal rate. This means a working model must also predict a dimensionless production rate that is greater than 1 for pro-high cells.

Since the dimensionless production rate of macrophage is always 1, a working model constrained by $meR^i_{RemRate}$ would yield that pro-high cells have greater PU.1 production rate than macrophages. Meanwhile, the model must also predict consistently with the measurement, where pro-high cells have lower PU.1 abundance than macrophage [1]. Together, a successful model requires that the PU.1 production rate is non-monotonic in response to PU.1 abundance.

However, in our model development (next section), we mainly rely on monotonic production rate functions such as Hill functions and statistical thermodynamic models. Given this choice of production rate functions, therefore, no models constrained by $meR^i_{RemRate}$ can predict pro-high correctly. To resolve this, we instead treated the 25th percentile of the $R^H_{RemRate}$ sample space as the removal rate constant. Because $R^H_{RemRate}$ decreases as $\tau_H$ increases, this lower removal rate constant reflects pro-high cell-cycle lengths in the upper portion of the $\tau_H$ distribution (around its 75th percentile). As shown in Figure S8, at this removal rate constant, production rate is allowed to be less than 1 (blue triangle marker), making it possible to find working models with a monotonic production rate function.

#### 2.4 Phenomenological models (Table S1)

We start by finding phenomenological model architecture that can reproduce the joint PU.1-GFP and synthesis-rate measurements of the "HSC differentiation dataset". Given that the removal rate constant of PU.1 is identical for all models, we include only the mathematical function for the PU.1 production rate from now on. For the model including an additional state variable, we include the full ODE for that particular variable as needed. For simplicity, even though all models are implemented in the dimensionless form, we focus on showing the dimensional equations. Equations of all phenomenological models are summarized in Table S1.

Table S1: Summary of phenomenological models. $i \in \{\text{B, L, H}\}$. For different models (Column 1), the production rate function of PU.1 in Eqn. (S24) is shown (Column 2). Column 3 shows the equation for the additional state variable. For the positive feedback model, in addition to the change in PU.1 abundance, the abundance of a generic TF, $[gTF]_i$, is also modeled (i.e., $Z_i = [gTF]_i$).

| | **Production**$([PU]_i, Z_i)$ | $\frac{d[gTF]_i}{dt}$ |
|---|---|---|
| AR&DGR | $k_{PU} \times \mathrm{h}([PU]_i)$ | N/A |
| Cell-type-specific autoregulation | $X_i \times \mathrm{h}([PU]_i)$ | N/A |
| Positive Feedback | $k_{feedback} \times \mathrm{h}([gTF]_i) \times \mathrm{h}([PU]_i)$ | $k_{gTF} \times \mathrm{h}([PU]_i) - \left(\frac{\log 2}{\tau_i} + D_{gTF}\right)[gTF]_i$ |

Hereafter, $\mathrm{h}(x)$ is the dimensionless Hill function (Eqn. (5) of the main text):

$$\mathrm{h}(x) = \tilde{b} + (1 - \tilde{b})\frac{x^N}{x^N + K^N} \tag{S27}$$

where $\tilde{b}$, $N$, and $K$ are the dimensionless basal level, the Hill coefficient, and the half-saturation constant, respectively. These parameters are specific to each Hill function: every occurrence of $\mathrm{h}(\cdot)$ in the models below has its own $\tilde{b}$, $N$, and $K$, even when two occurrences share the same input. The output of $\mathrm{h}(x)$ does not exceed 1; in each model, $\mathrm{h}(x)$ is multiplied by a scaling factor (e.g., $k_{PU}$) to be consistent with the unit of the corresponding equation.

##### 2.4.1 AR&DGR

In the "AR&DGR model", all three cell types share the same autoregulatory production rate function, i.e., regardless of the cell type, parameters of $k_{PU} \times \mathrm{h}([PU]_i)$ are identical. As such, the dependence of PU.1 production rate on PU.1 abundance is identical. Even though the solved steady-state $[PU]_i$ is cell-type-specific, the model turns out to be limited in producing steady states that agree with experimental data.

##### 2.4.2 Cell-type-specific autoregulation

We introduce a generic transcription factor, or $gTF$. Assuming the abundance of $gTF$ is cell-type-specific, some generic function of $gTF$, or $f([gTF])$, is then also cell-type-specific. As such, when the autoregulatory production rate function of PU.1, $\mathrm{h}([PU]_i)$, is multiplied by $f([gTF])$, the dependence of the overall PU.1 production rate on PU.1 abundance is now unique to each cell type. This is the "cell-type-specific autoregulation" model. In the simplest case, $f([gTF])$ can be some constant that is different for each cell type, $X_i$.

##### 2.4.3 Positive Feedback

Fitted $X_i$ from above leads us to consider a parsimonious realization where PU.1 and the generic TF are in mutual activation, as shown in the last row of Table S1. In addition to gTF activation on PU.1 production rate, gTF is now monotonically produced at a rate dependent on PU.1 abundance. The Hill parameters of h($[gTF]_i$) and h($[PU]_i$) are assumed to be distinct and independent.

Notably, the generic TF is removed with the same mechanism as PU.1, including both dilution ($\frac{\log 2}{\tau_i}[gTF]_i$) and degradation ($D_{gTF}[gTF]_i$), where $D_{gTF}$ is the natural degradation rate constant of gTF.

In the dimensionless model, the removal rate constant for gTF is $\frac{\frac{\log 2}{\tau_i}+D_{gTF}}{\frac{\log 2}{\tau_M}+D_{gTF}}$. Due to the unknown degradation rate constant $D_{gTF}$, error propagation in the macrophage-normalized removal rate constant is not possible, i.e., the error propagation method introduced in Section 1 does not apply to $\frac{\frac{\log 2}{\tau_i}+D_{gTF}}{\frac{\log 2}{\tau_M}+D_{gTF}}$. Therefore, we directly use the means of the cell-cycle length distributions constructed in Section 1.1 for $\tau_i$ and $\tau_M$ in the expression $\frac{\frac{\log 2}{\tau_i}+D_{gTF}}{\frac{\log 2}{\tau_M}+D_{gTF}}$. The only exception is pro-high cells, for which the 75th percentile of the $\tau_H$ distribution is used instead of its mean. This yields a lower gTF removal rate constant for pro-high cells, consistent with the choice made for PU.1 (Section 2.3).

#### 2.5 Biophysical models (Table S2)

In this section, we first broadly compare all attempted biophysical models. Then, we introduce the statistical thermodynamics framework used to compute the production rate in biophysical models. Subsequently, we explain how hypotheses of transcriptional regulation are implemented in the model. Lastly, we include the specific assumptions of the model.

To start with, equations of all biophysical models are summarized in Table S2.

##### 2.5.1 Models with C/EBP$\alpha$

1. C/EBP$\alpha$ model (without C/EBP$\alpha$ autoregulation)

   Literature search led us to conclude that C/EBP$\alpha$ is the second TF in feedback with PU.1. Moreover, on the upstream regulatory element (URE) of the PU.1 promoter, PU.1 and C/EBP$\alpha$ can bind to different DNA regions that are 14, 12, and 10 kilobase pairs away from the promoter ($-14$, $-12$, and $-10$ kb hereafter). Perturbations of PU.1 and C/EBP$\alpha$ binding sites on these DNA regions affect PU.1 gene expression [3]. We refer to the PU.1 expression measurements related to these perturbations as "PU.1 transcriptional dataset". Types of perturbation and cells in which the measurement was conducted are summarized in Table S6.

   Based on our previous work, a biophysical framework based on statistical mechanics can explain the patterns in the wildtype and perturbed gene expression measurements. This framework maps the abundance of transcription factors (TFs) regulating the gene to the transcription rate. Here, given that PU.1 and C/EBP$\alpha$ binding affect the PU.1 expression, we develop a biophysical function called Biophysical($[PU]_i, [CE]_i$) that outputs PU.1 transcription rate dependent on PU.1 and C/EBP$\alpha$ abundance.

   Separately, we assume that the wildtype gene expression is reasonable for estimating protein production rate. As such, the same function Biophysical($[PU]_i, [CE]_i$) not only explains the expression patterns in the "PU.1

Table S2: Summary of biophysical model equations, with $i \in \{B, L, H\}$. For different models (Column 1), Column 2 shows the production rate function of PU.1 in Eqn. (S24) and Column 3 shows the full ODE for the rate of change in C/EBP$\alpha$. The biophysical models mentioned in the main text are highlighted in gray.

| | **Production**($[PU]_i, Z_i$) | $\frac{d[CE]_i}{dt}$ |
|---|---|---|
| C/EBP$\alpha$ model (without C/EBP$\alpha$ autoregulation) | Biophysical($[PU]_i, [CE]_i$) | $k_{CE1} \times \mathrm{h}([PU]_i) - \left(\frac{\log 2}{\tau_i} + D_{CE}\right)[CE]_i$ |
| C/EBP$\alpha$ model (with C/EBP$\alpha$ autoregulation) | Biophysical($[PU]_i, [CE]_i$) | $k_{CE2} \times \mathrm{h}([PU]_i) \times \mathrm{h}([CE]_i) - \left(\frac{\log 2}{\tau_i} + D_{CE}\right)[CE]_i$ |
| C/EBP$\alpha$ model (open-loop) | Biophysical($[PU]_i, [CE]_i$) | N/A |
| RUNX1 model (without C/EBP$\alpha$ autoregulation) | Biophysical($[PU]_i, [CE]_i, [RUN]_i$) | $k_{CE1} \times \mathrm{h}([PU]_i) - \left(\frac{\log 2}{\tau_i} + D_{CE}\right)[CE]_i$ |
| RUNX1 model (with C/EBP$\alpha$ autoregulation) | Biophysical($[PU]_i, [CE]_i, [RUN]_i$) | $k_{CE2} \times \mathrm{h}([PU]_i) \times \mathrm{h}([CE]_i) - \left(\frac{\log 2}{\tau_i} + D_{CE}\right)[CE]_i$ |
| RUNX1 model (with RUNX1 activation) | Biophysical($[PU]_i, [CE]_i, [RUN]_i$) | $k_{CE3} \times \mathrm{h}([PU]_i) \times \mathrm{h}([RUN]_i) - \left(\frac{\log 2}{\tau_i} + D_{CE}\right)[CE]_i$ |

transcriptional dataset", but also serves as Production($[PU]_i, Z_i$) in Eqn. (S24), with $Z_i = [CE]_i$. Notably, **we assume the gene composition in "Prom141210" (see Table S6) is the wildtype DNA composition giving rise to the wildtype PU.1 expression.** Due to the dependence on $[CE]_i$, Biophysical($[PU]_i, [CE]_i$) takes care of both PU.1 autoregulation and activation from C/EBP$\alpha$.

On the other hand, the PU.1 protein removal rates are, as before, precomputed and fed into the model. Additionally, the remaining arm of the positive feedback, i.e., activation of C/EBP$\alpha$ by PU.1, is modeled with a Hill function, $k_{CE1} \times \mathrm{h}([PU]_i)$, due to the lack of gene expression data.

2. C/EBP$\alpha$ model (with C/EBP$\alpha$ autoregulation)

   Given that the model above failed, we considered adding the autoregulation of C/EBP$\alpha$. This is based on the experimental finding that there are C/EBP$\alpha$ binding sites on the +37 kb enhancer of the *Cebpa* gene [4]. Such an autoregulation is implemented with a phenomenological Hill function, $\mathrm{h}([CE]_i)$. This model failed to reproduce the joint data of PU.1-GFP and synthesis rates.

3. C/EBP$\alpha$ model (open-loop)
   Given that both models above failed, this model is one-dimensional: it removes the activation of C/EBP$\alpha$ by PU.1 and serves as a diagnostic tool. We treat abundances of C/EBP$\alpha$, or $[CE]_i$ in the production of PU.1, as unknown parameters to be optimized.

For all models in which C/EBP$\alpha$ is a state variable (in this section and in Section 2.5.2), the dimensionless C/EBP$\alpha$ removal rate constant, $\frac{\frac{\log 2}{\tau_i}+D_{CE}}{\frac{\log 2}{\tau_M}+D_{CE}}$, is precomputed in exactly the same way as that of PU.1. That is, it is obtained using the normalization and error-propagation procedure in Section 2.2, with $D_{PU}$ in Eqn. (S26) replaced by $D_{CE} = 0.4159\,\mathrm{h}^{-1}$ (calculated from a reported half-life of ~100 minutes; Table S8). As for PU.1, the mean of each resulting distribution is fed into the model, except for pro-high cells, for which the 25th percentile is used (Section 2.3).

##### 2.5.2 Models with C/EBP$\alpha$ and RUNX1

1. RUNX1 model (without C/EBP$\alpha$ autoregulation)
   The previous model cannot capture the experimental trends in cell-type-specific PU.1 abundances. To this end, we develop another model that additionally considers the effect of RUNX1, based on the observation that perturbations of RUNX1 binding sites on the same PU.1 URE affect PU.1 expression [5]. To incorporate this finding, we expand on the previous biophysical model such that RUNX1 regulates PU.1 expression. The new production rate function, Biophysical($[PU]_i, [CE]_i, [RUN]_i$), has dependence on RUNX1 abundance (i.e., $Z_i = ([CE]_i, [RUN]_i)$). This model is expected to explain the "expanded PU.1 transcriptional dataset", which consists of the "PU.1 transcriptional dataset" mentioned above, along with the newly incorporated measurements due to RUNX1 mutation [3, 5].

2. RUNX1 model (with C/EBP$\alpha$ autoregulation)
   Assuming low cooperativity of PU.1-dependent C/EBP$\alpha$ activation, adding the RUNX1 input does not improve the model's ability to explain the experimental data. To this end, we again include the C/EBP$\alpha$ autoregulation using the same phenomenological function as in Section 2.5.1. This model fitted well.

3. RUNX1 model (with RUNX1 activation)
   We also examine an alternative hypothesis that RUNX1 affects the production rate of C/EBP$\alpha$, in accordance with the reported RUNX1 binding sites on the C/EBP$\alpha$ enhancer [4]. The activation of C/EBP$\alpha$ by RUNX1 is implemented with a Hill function h($[RUN]_i$). This model did not fit well.

##### 2.5.3 A statistical-thermodynamics-based framework to model transcription rate

The functions Biophysical(...) are formulated to output transcription rate that corresponds to the experimental gene expression. To model transcription rate, we approximate it with the probability of RNAP being bound to DNA. The broad framework to compute RNAP binding probability follows Section (II.1) of the Supplementary Materials in [6]. Below, we expand on the previously developed framework [6] to model gene expression additionally affected by the presence of nucleosomes.

For a DNA molecule with T number of TF-binding sites and a single promoter, we assume each TF binding site can be bound (by either nucleosomes or TF), or unbound. Similarly, the promoter can be bound (by either nucleosomes or RNAP) or unbound. The binding configuration of the DNA molecule, $\alpha$, can be represented as:

$$\alpha = [\sigma_1, \sigma_2, ..., \sigma_T, \sigma_{pro}], \tag{S28}$$

where, $\sigma_k \in [0(unbound), 1(bound)]$ for $k = 1, 2, 3, \ldots, T$. Additionally, $\sigma_{pro} \in [0(unbound), 1(bound)]$. How we distinguish whether a specific site $\sigma$ is bound by TF or nucleosomes will be elucidated in a later section. We also define a specific set of $\alpha$ called $\alpha_T$. In this set, $\sigma_{pro} = 1$, denoting configurations with promoter being occupied by the RNAP.

The probability of DNA in configuration $\alpha$, in accordance with statistical mechanics, is defined as:

$$\text{Prob}_\alpha = \frac{\exp(-G_\alpha)}{\sum_\alpha \exp(-G_\alpha)} \tag{S29}$$

where, $G_\alpha$ is the free energy for the configuration $\alpha$, with unit $k_BT$:

$$G_\alpha = \sum_k \sigma_k (G_k^\circ - N_k \times \ln[L_k]) + \sum_{k,j,k<j} \sigma_k \sigma_j G_{kj}^\circ, \tag{S30}$$

with $k \in [1, 2, 3..., T, pro], j \in [1, 2, 3..., T, pro]$.

Specifically, $\sigma_k$ is the binary variable in Eqn. (S28) that denotes whether the site k is bound or unbound, $G^\circ_k$ is the standard binding energy when site $k$ is bound. $[L_k]$ is the concentration of the ligand (in our case, this can be TF, nucleosome, or RNAP) bound at site $k$. The variable $N_k$ is the ligand cooperativity at the particular site $k$. Additionally, $G^\circ_{kj}$ is the standard interaction energy when both site k and j are bound. Due to the way that $\sigma$ is incorporated, $G_\alpha$ only includes binding energies, interaction energies for two sites, as well as the abundances of ligands for those **occupied** sites.

For notation purposes, we let all energy parameters needed for Eqn. (S30) as the set $\{G^\circ\}$. Similarly, we define $\{[L]\}$ as the set of ligand abundances. Given individual configuration's probability, the RNAP binding probability, or the probability of transcription ("$\text{Prob}_{\text{trc}}$"), is the sum of probability over configurations with RNAP bound (i.e., the numerator is only contributed by configurations from the set $\alpha_T$)

$$\text{Prob}_{\text{trc}}(\{G^\circ\}, \{[L]\}) = \frac{\sum_{\alpha \in \alpha_T} \exp(-G_\alpha)}{\sum_\alpha \exp(-G_\alpha)} \tag{S31}$$

Note that the parentheses signify that this probability is a function of free energy parameters and ligand abundances. In our model, if the binding site $k$ is occupied by nucleosome or RNAP, $[L_k] = 1$. Otherwise, $[L_k]$ is equal to the corresponding transcription factor abundance.

With this theoretical framework, the cell-type-specific biophysical function Biophysical(...) listed in Table S2 is assumed to be equal to the RNAP binding probability. For example:

$$\text{Biophysical}([PU]_i, [CE]_i) = \text{Prob}_{\text{trc}}(\{G^\circ\}, \{[L]\}) \tag{S32}$$

where, in the set $\{[L]\}$, for sites $k$ that are PU.1 and C/EBP$\alpha$ binding sites, $[L_k]$ is equal to $[PU]_i$ and $[CE]_i$, respectively. Similarly, for the "RUNX1" models, $\text{Biophysical}([PU]_i, [CE]_i, [RUN]_i) = \text{Prob}_{\text{trc}}(\{G^\circ\}, \{[L]\})$. In the set $\{[L]\}$, for sites $k$ that are RUNX1 binding sites, $[L_k]$ is equal to $[RUN]_i$.

We would like to note two interesting points about this framework:

1. Connections to the Linear Framework
   Our model is equivalent to the "Linear Framework" at steady state and thermal equilibrium [7]. For example, for DNA with a single promoter, a reaction network contains only one reaction:

$$DNA + RNAP \xrightarrow{K_{eq}^{RNAP}} DNA : RNAP \tag{S33}$$

   Here, RNAP binds to DNA to form $DNA : RNAP$, with $K_{eq}^{RNAP}$ being the equilibrium constant.

   With the assumption that the ligand abundance (i.e., [RNAP]) is much larger than the abundance of DNA, the reaction can be estimated as

$$DNA \xrightarrow{K_{eq}^{RNAP}[RNAP]} DNA : RNAP \tag{S34}$$

   In this reaction, there are two possible states of DNA: unbound DNA or DNA:RNAP. As such, we assume the normalization condition that the total probability of finding DNA is 1.

When detailed balance is enforced, the steady state probability (see [7]) for the chemical species $\alpha$ is exactly $\text{Prob}_\alpha$ in Eqn. (S29). In this case, there are only two configurations ($\alpha$'s) considered by our model: $[\sigma_{pro} = 1]$ and $[\sigma_{pro} = 0]$. The steady-state probability of DNA:RNAP is equivalent to computing $\text{Prob}_\alpha$ for $\alpha = [\sigma_{pro} = 1]$. Here, $G_\alpha = G^\circ_{RNAP} - \ln[RNAP]$ in accordance with Eqn. (S30). Notably, comparing the statistical mechanics model to the steady state solution of the linear framework, it is observed that $G^\circ_{RNAP} = -\ln(K^{RNAP}_{eq})$.

2. The cooperativity term in Eqn. (S30)
   The cooperativity term $N_k$ takes an effect if a multimer form of the ligand is bound. For instance, consider an alternative reaction in the same reaction network,

$$DNA + 2RNAP \xrightarrow{K^{RNAP}_{eq}} DNA : 2RNAP \tag{S35}$$

   where now a dimer form of RNAP binds to the DNA. To compute the probability of finding $DNA : 2RNAP$, our model computes the probability of the configuration $\alpha = [\sigma_{pro} = 1]$. The energy landscape parameter for this configuration is now $G_\alpha = G^\circ_{RNAP} - 2 \times \ln([RNAP])$. It can be seen that the cooperativity of RNAP is reflected in front of the concentration term. In our model, for sites $k$ that are occupied by nucleosome or RNAP, $N_k = 1$. For sites $k$ that are occupied by PU.1, C/EBP$\alpha$, or RUNX1, $N_k$ is constrained by literature value ($N_k = 1$ if a monomer binds, $= 2$ if a dimer binds, etc.).

##### 2.5.4 Two mechanisms of gene regulation

Two types of transcriptional regulation are implemented for our models.

1. Thermodynamic control
   The first is thermodynamic control (details see section (IV.1.a) of [6]). The basic idea is that $G^\circ_{kj}$, the interaction energy, is nonzero if the two bound sites k and j are a combination of a TF-binding site and a promoter. That is, $G^\circ_{kj} \neq 0$ for $\sigma_k = 1$ and $\sigma_j = 1$, where $k \in [1, 2, 3 ... T]$ and $j = pro$.

2. Nucleosome destabilization
   The second mechanism, called "nucleosome destabilization", is directly motivated by a recently published paper [8]. Using single-molecule footprinting, the study showed that synthetic gene constructs harboring multiple TF-binding sites and a minimal promoter can be occupied by both TFs and nucleosomes. In particular, bound TF can recruit coactivators to destabilize nucleosome formation, increasing overall gene expression.

   For TF to activate gene expression via "nucleosome destabilization", we first refine our definition of the configuration $\alpha$ even further. For individual site $\sigma$ in $\alpha$, $\sigma_k \in [0(unbound), 1(TFbound), 1^*(nuc)]$ for $k = 1, 2, 3, \ldots, T$. Additionally, $\sigma_{pro} \in [0(unbound), 1(RNAPbound), 1^*(nuc)]$. The star(*) is to distinguish between TF and nucleosome bound at a particular site. During implementation, however, $\sigma$ can still only take values of 0 or 1, which determines whether a particular site contributes to $G_\alpha$.

   We introduce a positive (repulsive) energy landscape parameter if one site is TF-bound and another site is nucleosomal. That is, $G^\circ_{kj} > 0$ for $\sigma_k = 1^*$ and $\sigma_j = 1$, where $k \in [1, 2, 3 ... T, pro]$ and $j \in [1, 2, 3 ... T]$. In other words, TF can destabilize nucleosomes bound at either another TF-binding site or the promoter. For a configuration with $a$ number of TF bound and $b$ number of nucleosomes bound, there would be $a \times b$ number of the repulsive $G^\circ_{kj}$.

###### 2.5.5 Detailed assumptions made for biophysical models (Tables S3–S5)

In our work, all biophysical models consider a total of four binding sites. Any DNA configuration ($\alpha$) in the model can be specified as $\alpha = [\sigma_1, \sigma_2, \sigma_3, \sigma_{pro}]$, with $\sigma_k \in \{0, 1, 1^*\}$, in accordance with Eqn. (S28). This assumption is made to be consistent with the observation that there are three TF binding sites, including $-14$ kb ($\sigma_1$), $-12$ kb ($\sigma_2$), and $-10$ kb ($\sigma_3$) of the PU.1 upstream regulatory enhancer, as well as one PU.1 promoter ($\sigma_{pro}$), on the "wildtype" gene construct "Prom141210" (also see Table S6).

In each table below (Tables S3–S5), we enumerate all possible ways that $\sigma_1$, $\sigma_2$, $\sigma_3$, and $\sigma_{pro}$ can be occupied, respectively. Then, for each site $\sigma_k$, we further distinguish between the types of ligand (nucleosome or TF) bound. Depending on the types of ligand, listed are ligand-specific standard binding energy ($G^\circ_k$), abundance ($L_k$), and cooperativity ($N_k$). These parameters are used to compute the term $G^\circ_k - N_k \times \ln(L_k)$ in Eqn. (S30), for each unique $k \in \{1, 2, 3, \text{pro}\}$.

Here are several notes about the tables below:

1. The subscript to the energy term such as "nuc","PU", "CE", "RUN", and "RNAP" is used to specify the type of ligand that contributes to the free energy. These refer to nucleosome, PU.1, C/EBP$\alpha$, RUNX1, and RNA polymerase, respectively.

2. For the $-14$ kb site, the model considers the possibility that more than one type of ligand can be bound. For these types of occupancies, we can apply Eqn. (S30) recursively on this site, as if the site itself is its own "model". In this case, interaction energy $G^\circ_{kj}$ must be considered. The subscript with two ligand names combined (such as "CEPU") denotes the interaction.

3. For RNAP and nucleosome, ligand concentration is assumed to be 1, i.e., the effect of concentration is normalized and absorbed entirely into the standard binding energy term.

4. To obtain all possible configurations ($\alpha$) of the model, one can take the Cartesian product of the three tables (using Table S4 for both sites 2 and 3). When taking this product, for those configurations with TF-nucleosome or TF-RNAP bound simultaneously, one must further take into account the interaction $G^\circ_{kj}$ in accordance with Eqn. (S30).

Table S3: For the $-14$ kb region of the PU.1 upstream regulatory enhancer, or site 1 in the model, listed are all possible specific ways that the region can be occupied. Note that the presence of RUNX1 increases the number of possible occupancies from models without RUNX1 to models with RUNX1.

| **Biophysical Model Name** | **$-14$ kb region (site 1)** | | | |
|---|---|---|---|---|
| | **Ligand** | $\boldsymbol{\sigma_1}$ | $\boldsymbol{N_k \times \ln(L_k)}$ | $\boldsymbol{G_k^\circ}$ |
| Models without RUNX1 | Nucleosome | $1^*$ | $1 \times \ln(1) = 0$ | $G_{nuc}^\circ$ |
| | Nothing | 0 | N/A | 0 |
| | C/EBP$\alpha$ alone | 1 | $2 \times \ln([CE]_i)$ | $G_{CE}^\circ$ |
| | PU.1 alone | 1 | $1 \times \ln([PU]_i)$ | $G_{PU}^\circ$ |
| | Both C/EBP$\alpha$ and PU.1 | 1 | $2 \times \ln([CE]_i) + 1 \times \ln([PU]_i)$ | $G_{CE}^\circ + G_{PU}^\circ + G_{CEPU}^\circ$ |
| Models with RUNX1 | Nucleosome | $1^*$ | $1 \times \ln(1) = 0$ | $G_{nuc}^\circ$ |
| | Nothing | 0 | N/A | 0 |
| | PU.1 alone | 1 | $1 \times \ln([PU]_i)$ | $G_{PU}^\circ$ |
| | C/EBP$\alpha$ alone | 1 | $2 \times \ln([CE]_i)$ | $G_{CE}^\circ$ |
| | RUNX1 alone | 1 | $2 \times \ln([RUN]_i)$ | $G_{RUN}^\circ$ |
| | Both PU.1 and C/EBP$\alpha$ | 1 | $1 \times \ln([PU]_i) + 2 \times \ln([CE]_i)$ | $G_{PU}^\circ + G_{CE}^\circ + G_{CEPU}^\circ$ |
| | Both PU.1 and RUNX1 | 1 | $1 \times \ln([PU]_i) + 2 \times \ln([RUN]_i)$ | $G_{PU}^\circ + G_{RUN}^\circ + G_{PURUN}^\circ$ |
| | Both C/EBP$\alpha$ and RUNX1 | 1 | $2 \times \ln([CE]_i) + 2 \times \ln([RUN]_i)$ | $G_{CE}^\circ + G_{RUN}^\circ + G_{CERUN}^\circ$ |
| | PU.1, C/EBP$\alpha$, and RUNX1 | 1 | $1 \times \ln([PU]_i) + 2 \times \ln([CE]_i) + 2 \times \ln([RUN]_i)$ | $G_{PU}^\circ + G_{CE}^\circ + G_{RUN}^\circ + + G_{CEPU}^\circ + G_{CERUN}^\circ + G_{PURUN}^\circ$ |

Table S4: For the $-12$ kb region, or site 2 in the model, listed are all possible specific ways that this region can be occupied. The possible occupancy of the $-10$ kb region, or site 3, is identical to site 2. There is no difference between models with or without RUNX1.

| **Biophysical Model Name** | **$-12$ kb or $-10$ kb region (site 2 or site 3)** | | | |
|---|---|---|---|---|
| | **Ligand** | $\boldsymbol{\sigma_2}$ **or** $\boldsymbol{\sigma_3}$ | $\boldsymbol{N_k \times \ln(L_k)}$ | $\boldsymbol{G_k^\circ}$ |
| All biophysical models | Nucleosome | $1^*$ | $1 \times \ln(1) = 0$ | $G_{nuc}^\circ$ |
| | Nothing | 0 | N/A | 0 |
| | PU.1 | 1 | $1 \times \ln([PU]_i)$ | $G_{PU}^\circ$ |

Table S5: For the promoter, or site "pro" in the model, listed are all possible specific ways that the region can be occupied. There is no difference between models with or without RUNX1.

| **Biophysical Model Name** | **Promoter (site pro)** | | | |
|---|---|---|---|---|
| | **Ligand** | $\boldsymbol{\sigma_{pro}}$ | $\boldsymbol{N_k \times \ln(L_k)}$ | $\boldsymbol{G_k^\circ}$ |
| All biophysical models | Nucleosome | $1^*$ | $1 \times \ln(1) = 0$ | $G_{nuc}^\circ$ |
| | Nothing | 0 | N/A | 0 |
| | RNAP | 1 | $1 \times \ln(1) = 0$ | $G_{RNAP}^\circ$ |

### 3 Further normalization of models to bypass unknown dimensional steady state

In Section 2, Eqn. (S21), the analysis of the dimensionless model is possible given that the steady state of the dimensional model, $[PU^*]_M$, is known. We obtained this value by replicating the AR&GF model [1]. However, in the dimensionless model we develop (Eqn. (S25)), the true steady state $[PU^*]_M$ remains unknown. Here, we demonstrate how to bypass this via further normalization of the model [2].

#### 3.1 Phenomenological models

Given that $[PU^*]_M$ only appears in the dimensionless production rate term, i.e., $\frac{\text{Production}([pu]_i[PU^*]_M, Z_i)}{\text{Production}([PU^*]_M, Z^*_M)}$, here we only need to focus on the Production(...) part of the model.

Let's use $\text{Production}([PU]_i, Z_i) = k_{PU} \times \text{h}([PU]_i)$ as an example. In this case, the scaling factor $k_{PU}$ cancels, and the dimensionless production rate is $\frac{\text{h}([pu]_i[PU^*]_M)}{\text{h}([PU^*]_M)}$. The numerator of the dimensionless rate is:

$$\text{h}([pu]_i[PU^*]_M) = \tilde{b} + (1-\tilde{b})\frac{([pu]_i[PU^*]_M)^N}{([pu]_i[PU^*]_M)^N + K^N} \tag{S36}$$

which is equivalent to

$$\text{h}([pu]_i[PU^*]_M) = \tilde{b} + (1-\tilde{b})\frac{([pu]_i)^N}{([pu]_i)^N + \left(\frac{K}{[PU^*]_M}\right)^N} \tag{S37}$$

The right-hand side is just a Hill function with input $[pu]_i$ in terms of a different half-saturation constant, $K/[PU^*]_M$.

Therefore, if we let $K' = K/[PU^*]_M$, we effectively transformed $\text{h}([pu]_i[PU^*]_M)$ in terms of $K$ into $\text{h}([pu]_i)$ in terms of $K'$, that is,

$$\text{h}([pu]_i[PU^*]_M, K) = \text{h}([pu]_i, K') \tag{S38}$$

On the other hand, for the expression in the denominator,

$$\text{h}([PU^*]_M) = \tilde{b} + (1-\tilde{b})\frac{([PU^*]_M)^N}{([PU^*]_M)^N + K^N} \tag{S39}$$

which can be further nondimensionalized and equivalently expressed as

$$\text{h}([PU^*]_M) = \tilde{b} + (1-\tilde{b})\frac{1^N}{1^N + \left(\frac{K}{[PU^*]_M}\right)^N} \tag{S40}$$

The right-hand side indicates that it is a Hill function with input 1 and half-saturation constant $K' = \frac{K}{[PU^*]_M}$. As such,

$$\text{h}([PU^*]_M, K) = \text{h}(1, K') \tag{S41}$$

Together, the dimensionless production rate now becomes the following:

$$\frac{\text{Production}([pu]_i[PU^*]_M, Z_i)}{\text{Production}([PU^*]_M, Z_M^*)} = \frac{\text{h}([pu]_i[PU^*]_M, K)}{\text{h}([PU^*]_M, K)} = \frac{\text{h}([pu]_i, K')}{\text{h}(1, K')} \tag{S42}$$

As such, we can directly simulate $\frac{d[pu]_i}{dt}$ without having to know $[PU^*]_M$. Fitting the model to find the parameter $K'$ already takes care of $[PU^*]_M$. On the other hand, the remaining Hill function parameters ($N$ and $\tilde{b}$) are unaffected by normalization.

#### 3.2 Biophysical models

A similar operation can be done for the biophysical model. Consider the case where Biophysical($[PU]_i$) has the form

$$\text{Biophysical}([PU]_i) = \text{Prob}_{\text{trc}}([PU]_i) = \frac{e^{-(G^\circ - \ln([PU]_i))}}{1 + e^{-(G^\circ - \ln([PU]_i))}} \tag{S43}$$

where the RNAP binding probability depends on the abundance of PU.1 in cell type $i$.

The numerator of the dimensionless production rate is then:

$$\text{Prob}_{\text{trc}}([pu]_i[PU^*]_M) = \frac{e^{-(G^\circ - \ln([pu]_i[PU^*]_M))}}{1 + e^{-(G^\circ - \ln([pu]_i[PU^*]_M))}} \tag{S44}$$

Here, we define $G'^\circ = G^\circ - \ln([PU^*]_M)$. As such, the expression above can be equivalently expressed as

$$\text{Prob}_{\text{trc}}([pu]_i[PU^*]_M) = \frac{e^{-(G'^\circ - \ln([pu]_i))}}{1 + e^{-(G'^\circ - \ln([pu]_i))}} \tag{S45}$$

where the right-hand side has the same form as $\text{Prob}_{\text{trc}}$ in terms of $[pu]_i$, just with a different energy $G'^\circ$.

By absorbing $[PU^*]_M$ into a newly defined energy term, we made the following transformation:

$$\text{Prob}_{\text{trc}}([PU]_i, G^\circ) = \text{Prob}_{\text{trc}}([pu]_i, G'^\circ) \tag{S46}$$

On the other hand, $\text{Prob}_{\text{trc}}([PU^*]_M, G^\circ)$ is just $\text{Prob}_{\text{trc}}([pu]_i = 1, G'^\circ)$, given that $[pu]_i = [PU]_i/[PU^*]_M$. Omitting the derivation, this can be obtained by explicitly writing out $\text{Prob}_{\text{trc}}([PU^*]_M, G^\circ)$ and substituting $G'^\circ = G^\circ - \ln([PU^*]_M)$.

As such,

$$\frac{\text{Production}([pu]_i[PU^*]_M, Z_i)}{\text{Production}([PU^*]_M, Z_M^*)} = \frac{\text{Prob}_{\text{trc}}([pu]_i[PU^*]_M, G^\circ)}{\text{Prob}_{\text{trc}}([PU^*]_M, G^\circ)} = \frac{\text{Prob}_{\text{trc}}([pu]_i, G'^\circ)}{\text{Prob}_{\text{trc}}(1, G'^\circ)} \tag{S47}$$

We need only solve for the energy parameter $G'^{\circ}$ without knowing the true steady state of the dimensional ODE. On the other hand, interaction energy terms $G^{\circ}_{kj}$ are unaffected by the normalization method.

### 4 Extract information on PU.1 cooperativity from the open-loop C/EBP$\alpha$ model

This section complements Results 3.4 and explains how we diagnosed insufficient cooperativity in the closed-loop C/EBP$\alpha$ model by using the open-loop version.

Given that the "C/EBP$\alpha$ model (open-loop)" is a one-dimensional ODE, whereas the "C/EBP$\alpha$ model (without C/EBP$\alpha$ autoregulation)" is two-dimensional, we first convert the latter into a one-dimensional model to seek an expression of C/EBP$\alpha$ that can relate the two models.

For the "C/EBP$\alpha$ model (without C/EBP$\alpha$ autoregulation)", the dimensionless governing equation for C/EBP$\alpha$ is as follows:

$$\frac{d[ce]_i}{dt} = \frac{\mathrm{h}([pu]_i[PU^*]_M)}{\mathrm{h}([PU^*]_M)} - \frac{\frac{\log 2}{\tau_i} + D_{CE}}{\frac{\log 2}{\tau_M} + D_{CE}}[ce]_i \tag{S48}$$

To eliminate the C/EBP$\alpha$ dimension, we can express the dimensionless C/EBP$\alpha$ abundance as a function of $[pu]_i$ only, using the steady state of $\frac{d[ce]_i}{dt}$. Let this steady state be $[ce^*]_i^{\text{feedback}}$, which can be computed using $\frac{d[ce]_i}{dt} = 0$:

$$[ce^*]_i^{\text{feedback}} = \frac{\frac{\mathrm{h}([pu]_i[PU^*]_M)}{\mathrm{h}([PU^*]_M)}}{\frac{\frac{\log 2}{\tau_i}+D_{CE}}{\frac{\log 2}{\tau_M}+D_{CE}}} = \frac{\frac{\mathrm{h}([pu]_i, K')}{\mathrm{h}(1, K')}}{\frac{\frac{\log 2}{\tau_i}+D_{CE}}{\frac{\log 2}{\tau_M}+D_{CE}}} \tag{S49}$$

Note that Eqns. (S48) and (S49) are written with explicit $\tau_i$, $\tau_M$, and $D_{CE}$ for derivation purposes only. In implementation, the error-propagated dimensionless C/EBP$\alpha$ removal rate constants (Section 2.5.1) are used.

Separately, let the freely-optimized dimensionless C/EBP$\alpha$ and PU.1 abundances from the "C/EBP$\alpha$ model (open-loop)" be $[ce]_i^{\text{no feedback}}$ and $[pu]_i^{\text{no feedback}}$.

When all other model parameters (i.e., energy landscape parameters in the PU.1 biophysical production rate function) are the same, then two models should be equivalent such that

$$[ce^*]_i^{\text{feedback}} = [ce]_i^{\text{no feedback}} \tag{S50}$$

Given the freely optimized $[ce]_i^{\text{no feedback}}$ and $[pu]_i^{\text{no feedback}}$, we can find Hill parameters in Eqn. (S49) such that $[ce^*]_i^{\text{feedback}}$ approaches the known values of $[ce]_i^{\text{no feedback}}$. All other parameters, such as C/EBP$\alpha$ degradation rate ($D_{CE}$) and cell-cycle length ($\tau$) in Eqn. (S49) are known.

### 5 Model fitting

In this section, we demonstrate fitting procedures. Since all models are formulated to reproduce the joint PU.1-GFP and synthesis-rate measurements of the "HSC differentiation dataset", we first show the formulation of objective functions for these measurements. Then, for biophysical models, we show additional modifications to the objective function that allow them to fit to transcriptional data. Lastly, we include a comparison for the parameter space of all models examined in our work, as well as a summary of experimental evidence used as model constraints.

Overall,

1. All models are implemented in MATLAB and fitting is done via MATLAB (2024a) particle swarm optimization [9].
2. For models without positive feedback, each model contains three ODEs describing the rate of change in PU.1 abundance ($\frac{d[pu]_B}{dt}$, $\frac{d[pu]_L}{dt}$, $\frac{d[pu]_H}{dt}$). Each cell-type-specific ODE differs from each other due to the removal rate. For models with feedback, there are three **systems** of ODEs, with each system containing two ODEs. The first ODE describes the rate of change in PU.1, and the second the rate of change in the generic TF or C/EBP$\alpha$.

#### 5.1 Fitting to the joint PU.1-GFP and synthesis-rate measurements

To fit models to the joint PU.1-GFP and synthesis-rate measurements of the "HSC differentiation dataset", the objective function is intended to minimize the difference between experimental PU.1-GFP abundance and model-predicted PU.1 abundance. It also minimizes the difference between the experimental PU.1 synthesis rate and model-predicted PU.1 production rate. The workflow can be dissected as such:

1. simulate ODEs, or systems of ODEs, to get a rough estimate of equilibrium point(s)
2. use the equilibrium points as "x0" for the MATLAB function $fsolve$ to solve for equilibrium point(s)
3. test the stability of $fsolve$ solutions by examining the eigenvalue(s). If real parts of all eigenvalues are negative, $fsolve$ solutions are sufficient. If not, use ODE solvers with tight tolerance, using $fsolve$ solutions as initial conditions
4. for each cell type, either the fsolve solution or the end point of the simulated state-variable trajectory is recorded and treated as the predicted dimensionless PU.1 stable steady state, $[pu^*]^i_{pred}$, with $i \in \{\mathrm{B, L, H}\}$.
5. concurrent with step 4, the dimensionless production rate evaluated at $[pu^*]^i_{pred}$ is recorded as $v^{*i}_{pred}$.
6. for each $[pu^*]^i_{pred}$ and $v^{*i}_{pred}$, apply the following penalty function:

$$\text{penalty}(lb, x, ub) = g1 \times \text{heaviside}(lb - x) \times (\frac{lb - x}{x})^2 + g2 \times \text{heaviside}(lb - x) \\ + g1 \times \text{heaviside}(x - ub) \times (\frac{x - ub}{x})^2 + g2 \times \text{heaviside}(x - ub) \quad \text{(S51)}$$

The function leads to a nonzero positive number when $x < lb$ or $x > ub$, so as to penalize the input $x$ to stay in the range [lb,ub]. Moreover, the bigger the difference is between $x$ and the bound, the larger the penalty. The constants g1 and g2 are assumed to be 1 and 0.1, respectively.

7. Given a distribution of error-propagated, normalized measurement $R^i_{mt}$, let the lower bound and the upper bound of the distribution be $R^i_{mt}|_{lb}$ and $R^i_{mt}|_{ub}$ (with the exception of $R^B_{GFP}|_{lb}$, which is set to approximately 1/200; see Section 1.2). The score, $S_{penalty}$, to be minimized by the objective function is defined as such:

$$S_{penalty} = \sum_{i \in \{\mathrm{B,L,H}\}} \text{penalty}(R^i_{GFP}|_{lb}, [pu^*]^i_{pred}, R^i_{GFP}|_{ub}) + \sum_{i \in \{\mathrm{B,L,H}\}} \text{penalty}(R^i_{syn}|_{lb}, v^{*i}_{pred}, R^i_{syn}|_{ub}) \quad \text{(S52)}$$

In other words, we enforce the model to predict PU.1 stable states and production rates at these stable states to stay in the corresponding error-propagated experimental range of PU.1-GFP and PU.1 synthesis rate, respectively.

#### 5.2 Fitting to "PU.1 transcriptional dataset" and "expanded PU.1 transcriptional dataset" (Table S6)

In addition to fitting to the joint PU.1-GFP and synthesis-rate measurements, the biophysical models are also fitted to PU.1 gene expression data. Let PU.1 expression measured from cell type $i$ for gene construct $gc$ be $F^i_{gc}$. Based on the available data, $i \in \{\mathrm{B, M}\}$ and $gc \in \{Prom, Prom10, Prom12, \ldots, BProm1412\}$. The full list of gene constructs is listed in Table S6.

Table S6: List of PU.1 gene constructs of which the expression is used to quantitatively constrain PU.1 production rate in the biophysical models. **Note, "PU.1 transcriptional dataset" refers to all data points except for "Prom14 (RUN14mut)". On the other hand, "expanded PU.1 transcriptional dataset" includes all datapoints.**

| Cell type | Data point label | Genetic perturbations | Reference |
|---|---|---|---|
| Macrophages | "Prom" | PU.1 promoter only | Figure 4D of [3] |
| | "Prom10" | −10 kb and PU.1 promoter | |
| | "Prom12" | −12 kb and PU.1 promoter | |
| | "Prom14" | −14 kb and PU.1 promoter | |
| | "Prom1410" | −14 kb, −10 kb, and PU.1 promoter | |
| | "Prom1412" | −14 kb, −12 kb, and PU.1 promoter | |
| | "Prom141210" | −14 kb, −12 kb, −10 kb, and PU.1 promoter | |
| | "Prom1412 (PU12mut)" | −14 kb, −12 kb, and PU.1 promoter, with PU.1 site at −12 kb mutated | Figure 6F of [3] |
| | "Prom1412 (CE14mut)" | −14 kb, −12 kb, and PU.1 promoter, with all three C/EBP$\alpha$ sites at −14 kb mutated | |
| | "Prom1412 (PU12CE14mut)" | −14 kb, −12 kb, and PU.1 promoter, with all three C/EBP$\alpha$ sites at −14 kb mutated and PU.1 site at −12 kb mutated | |
| | "Prom14 (RUN14mut)" | −14 kb and PU.1 promoter, with all three AML1 sites at −14 kb mutated | Figure 1D of [5] |
| B cells | "BProm" | PU.1 promoter only | Figure 4E of [3] |
| | "BProm14" | −14 kb and PU.1 promoter | |
| | "BProm1412" | −14 kb, −12 kb, and PU.1 promoter | |

##### 5.2.1 Processing of transcriptional data

To feed all transcriptional data into the model, additional data processing is conducted to make sure they are on the same scale.

In [3], luciferase reporter activity in macrophages and B cells was reported with respect to the promoter-only construct in the same cell type (Figures 4D and 4E of [3]). That is, we know how PU.1 reporter activity compares between perturbations in the same cell type. But it is unclear how activity compares between cell types. To this end, we introduce an unknown parameter called "cell-type rescaling factor" ($\varepsilon$) such that

$$\frac{F^{B}_{BProm}}{F^{M}_{Prom}} = \varepsilon \tag{S53}$$

where, "BProm" is the data point with a gene construct with PU.1 promoter alone measured in B cells, and "Prom" is the construct with PU.1 promoter alone measured in macrophages. With this factor, we can express all measurements in B cells to be in terms of the promoter-only PU.1 activity in macrophage (i.e., "Prom"). The parameter $\varepsilon$ is fitted.

Additionally, the measurement provided in Figure 6F of [3] was reported as relative percentage with respect to the data measured from a gene construct with promoter, −12 kb, and −14 kb. We assume the activity of this construct (having 100% expression) is the same as the construct with promoter, −12 kb, and −14 kb shown in Figure 4D [3]. This way, we can express the expression of constructs "Prom1412 (PU12mut)", "Prom1412 (CE14mut)", and "Prom1412 (PU12CE14mut)" using reporter activity, with respect to the construct "Prom".

In a separate paper reporting PU.1 expression upon RUNX1 site mutation [5], the study measured the luciferase activity for the construct with −14 kb and promoter, before and after RUNX1 binding site mutation. To reconcile reporter activity measured from different experimental conditions between papers, we assume this construct without RUNX1 mutation in [5], has the same expression as the "Prom14" in [3]. This way, the RUNX1-mutated expression can then be converted to be on the same scale as the remaining data points.

Data processing leads to all data points being expressed as relative fold changes with respect to the "Prom" data point. We further normalize all data by the largest measurement ("Prom1412") such that the experimental data is now between 0 and 1, directly corresponding to the probability computed by the biophysical function.

##### 5.2.2 Using steady-state PU.1 abundance constrained by the joint PU.1-GFP and synthesis-rate measurements to fit to gene expression data

Using previously formulated biophysical models, we first define $\text{Prob}^{gc}_{\text{trc}}$ as the RNAP binding probability computed for gene construct $gc$. Then, the cell-type-specific ODE systems to be simulated in **Step 1 of Section 5.1** are:

$$\frac{d[pu]_i}{dt} = \frac{\text{Prob}^{Prom141210}_{\text{trc}}([pu]_i, [tf1]_i, [tf2]_i, \ldots, \{G'^{\circ}\})}{\text{Prob}^{Prom141210}_{\text{trc}}(1, 1, 1, \ldots, \{G'^{\circ}\})} - \frac{\frac{\log 2}{\tau_i} + D_{PU}}{\frac{\log 2}{\tau_M} + D_{PU}}[pu]_i \tag{S54}$$

where $\text{Prob}^{Prom141210}_{\text{trc}}$ is the RNAP binding probability for the gene construct giving rise to "Prom141210" (assumed to be wildtype). Meanwhile, $[tf1]_i, [tf2]_i, \ldots$ are dimensionless abundances of additional non-PU.1 TFs (i.e., the elements of $Z_i$) that the biophysical function depends on, based on the specific models listed in Table S2. This reflects our assumption that, regardless of cell type, steady state PU.1 abundance occurs in the wildtype PU.1 construct.

Next, as before, the steady state found by the ODE solver, $[pu^*]^i_{pred}$, and RNAP binding probability at this steady state, are subjected to the penalty in accordance with Eqn. (S51):

$$S_{penalty1} = \sum_{i\in\{\text{B,L,H}\}} \text{penalty}\left(R^i_{GFP}|_{lb}, [pu^*]^i_{pred}, R^i_{GFP}|_{ub}\right) + \sum_{i\in\{\text{B,L,H}\}} \text{penalty}\left(R^i_{syn}|_{lb}, \frac{\text{Prob}^{Prom141210}_{\text{trc}}([pu^*]^i_{pred}, [tf1^*]^i_{pred}, [tf2^*]^i_{pred}, \ldots, \{G'^\circ\})}{\text{Prob}^{Prom141210}_{\text{trc}}(1,1,1,\ldots,\{G'^\circ\})}, R^i_{syn}|_{ub}\right) \tag{S55}$$

At this point, the solver has found steady states $[pu^*]^i_{pred}$. If the model is a system of ODEs, it has also found $[tf1^*]^i_{pred}$, $[tf2^*]^i_{pred}$, etc. Given these steady states, the objective function can compute RNAP binding probability for all constructs in the "PU.1 transcriptional dataset" or "expanded PU.1 transcriptional dataset" (listed in Table S6). To do so, for each $F^i_{gc}$, the function predicts a corresponding RNAP binding probability using the function

$$\text{Prob}^{gc}_{\text{trc}}([pu^*]^i_{pred}, [tf1^*]^i_{pred}, [tf2^*]^i_{pred}, \ldots, \{G'^\circ\}) \tag{S56}$$

i.e., under the same set of energy landscape parameters ($\{G'^\circ\}$), the objective function computes RNAP binding probability for different gene constructs ($gc$) in different cell types ($i$).

The difference between $gc$ is taken care of by the number of configurations used to compute $\text{Prob}^{gc}_{\text{trc}}$. The difference in cell types is reflected by using solver-found steady-state unique to that cell type. Specifically,

$$[pu^*]^i_{pred} = \begin{cases} 1 & \text{for } i = M \\ [pu^*]^B_{pred} & \text{for } i = B \end{cases} \tag{S57}$$

Similarly,

$$[tf^*]^i_{pred} = \begin{cases} 1 & \text{for } i = M \\ [tf^*]^B_{pred} & \text{for } i = B \end{cases} \tag{S58}$$

With experimental reporter activity, or $F^i_{gc}$, and PU.1 synthesis rate computed from Eqn. (S56), the biophysical model minimizes the following normalized mean square error:

$$S_{mse} = \frac{1}{n} \sum_{i\in\{\text{B,M}\}} \sum_{gc\in\{Prom, Prom10, Prom12, \ldots, BProm1412\}} \frac{\left(\text{Prob}^{gc}_{\text{trc}}([pu^*]^i_{pred}, [tf1^*]^i_{pred}, [tf2^*]^i_{pred}, \ldots, \{G'^\circ\}) - F^i_{gc}\right)^2}{\left(F^i_{gc}\right)^2} \tag{S59}$$

where $n$ is the number of data points.

##### 5.2.3 Using RNA-seq data to constrain RUNX1 abundances and evaluate C/EBP$\alpha$ production rate

Lastly, we introduce the constraints imposed on RUNX1 in biophysical models, as well as how we evaluate the predicted C/EBP$\alpha$ production rate. The RNA-seq measurements for the *Cebpa* and *Runx1* genes are obtained from Haemosphere [10]. Let the raw RNA-seq measurement for the gene ($gn$) and cell lineages ($cl$) be $A^{cl}_{gn}$. Specifically, $cl \in \{\text{Multi Potential Progenitor, B Cell Lineage, Macrophage Lineage}\}$ and $gn \in \{Cebpa, Runx1\}$. Notably, the Haemosphere measurements for progenitor cells do not distinguish between pro-low and pro-high cells for our project.

All RNA-seq data are reported in the unit of $\log_2(\text{TPM}+1)$ (TPM = "Transcripts Per Million"; addition of 1 prevents the expression from being undefined when the expression is zero). To estimate the **ratios** in measurement between cell types in the unit of TPM, the following transformation on raw data is conducted. For a particular gene ($gn$) in a certain cell lineage ($cl$), given that the TPM values for most lineages are much greater than 1, the RNA-seq measurement normalized by macrophage ("$normA_{gn}^{cl}$") is approximated as:

$$normA_{gn}^{cl} \approx 2^{(A_{gn}^{cl} - A_{gn}^{\text{Macrophage Lineage}})} \tag{S60}$$

Within each cell lineage $cl$, the measurements vary significantly due to different experimental conditions. We record the minimum and maximum measurements $A_{gn}^{cl}|_{min}$ and $A_{gn}^{cl}|_{max}$, respectively. Given the maximum and minimum measurement in each cell lineage, we then compute the macrophage-normalized minimum and maximum RNA-seq measurement for a particular cell lineage ($cl$) for a particular gene ($gn$) as

$$normA_{gn}^{cl}|_{max} \approx 2^{(A_{gn}^{cl}|_{max} - A_{gn}^{\text{Macrophage Lineage}}|_{min})} \tag{S61}$$

and

$$normA_{gn}^{cl}|_{min} \approx 2^{(A_{gn}^{cl}|_{min} - A_{gn}^{\text{Macrophage Lineage}}|_{max})} \tag{S62}$$

**For RUNX1** Our models do not compute RUNX1 production rate; instead, its calculations require the abundances of RUNX1. To constrain the RUNX1 abundances across cell types, we decide to use RNA-seq data, an estimate for protein production rate. This assumption is due to the extremely fast protein degradation rate.

Suppose we were to model RUNX1 abundance using the same principles as Eqn. (S25):

$$\frac{d[run]_i}{dt} = \frac{\text{Production}([run]_i[RUN^*]_M)}{\text{Production}([RUN^*]_M)} - \frac{\frac{\log 2}{\tau_i} + D_{RUN}}{\frac{\log 2}{\tau_M} + D_{RUN}}[run]_i \tag{S63}$$

where $[RUN]$, $[run]$, and $D_{RUN}$ are dimensional abundance, dimensionless abundance, and natural degradation rate constant of RUNX1. As reported in literature, the half-life of RUNX1 is ~60 min, or 1 hour [11], leading to $D_{RUN} = \log(2)/(1\,\text{h}) = \log(2)\,\text{h}^{-1}$. On the other hand, the minimum cell-cycle length $\tau_i$ across cell types is ~11 hours (pro-high cells). Since the protein decays much faster than being diluted upon cell division, the dimensionless removal rate constant, or $\frac{\frac{\log 2}{\tau_i}+D_{RUN}}{\frac{\log 2}{\tau_M}+D_{RUN}}$, approaches 1.

As such, according to Eqn. (S63), at steady state ($\frac{d[run]_i}{dt} = 0$),

$$\frac{\text{Production}([run]_i[RUN^*]_M)}{\text{Production}([RUN^*]_M)} = [run]_i \tag{S64}$$

In other words, steady-state RUNX1 abundance approaches its production rate.

In line with this, we constrain the dimensionless RUNX1 abundance with macrophage-normalized RNA-seq data:

$$S_{penalty2} = \sum_{i \in \{\mathrm{B,L,H}\}} \mathrm{penalty}(normA^{cl(i)}_{Runx1}|_{min}, [run]_i, normA^{cl(i)}_{Runx1}|_{max}) \tag{S65}$$

Here, the cell lineage $cl(i)$ corresponding to each cell type $i$ is

$$\begin{cases} cl = \text{Multi Potential Progenitor} & \text{if } i = L \\ cl = \text{Multi Potential Progenitor} & \text{if } i = H \\ cl = \text{B Cell Lineage} & \text{if } i = B \end{cases} \tag{S66}$$

Together, for biophysical models, the error to be minimized ($S$) is the sum of Eqns. (S55), (S59), and (S65):

$$S = S_{penalty1} + S_{penalty2} + S_{mse} \tag{S67}$$

**For C/EBP$\alpha$** On the other hand, we directly evaluate predicted C/EBP$\alpha$ production rate using the $normA^{cl}_{Cebpa}|_{min}$ and $normA^{cl}_{Cebpa}|_{max}$, with the same mapping between model prediction (specified by $i$) and RNA-seq data (specified by $cl$) for cell types, as in Eqn. (S66). Notably, we decided not to constrain C/EBP$\alpha$ production rate using RNA-seq data during model fitting, since all models failed to fit to C/EBP$\alpha$ RNA-seq data. This suggests the constraints on C/EBP$\alpha$ production rate are not helpful in identifying a working model.

##### 5.3 Summary of all model parameters and parameter constraints (Tables S7 and S8)

For reference, the constrained and fitted parameters for each model are summarized in Table S7. In Table S8, we list experimental data that constrain biophysical model parameters.

Table S7: Summary of model parameters, with $i \in \{B, L, H\}$ and "parameters" abbreviated as "pars". For different models in our study (Column 1), shown are the different type(s) of datasets used for fitting (Column 2), constrained parameters (Column 3), and unknown parameters to be fitted using optimization (Column 4). For each Hill function, the "Hill parameters" refer to its scaling factor (e.g., $k_{PU}$), $\tilde{b}$, $N$, and $K'$ (dimensionless due to the normalization shown in Section 3). Similarly, $\{G'^{\circ}\}$ denotes the energy parameters of the biophysical functions after the normalization shown in Section 3. The parameter $\varepsilon$ ("cell-type rescaling factor") is discussed in Section 5.2.1. Also, for all models, the removal rate constant is constrained/precomputed (see Eqn. (S26)). For simplicity, they are not listed in Column 3. In Column 2, "HSC differentiation dataset" refers to its joint PU.1-GFP and synthesis-rate measurements. Models discussed in the main text are colored in gray.

| **Model** | **Data** | **Constrained pars** | **Fitted pars** |
|---|---|---|---|
| AR&DGR | HSC differentiation dataset | N/A | • Hill pars of h($[PU]_i$) |
| Cell-type-specific autoregulation | HSC differentiation dataset | N/A | • Hill pars of h($[PU]_i$)<br>• $X_i$ |
| Positive Feedback | HSC differentiation dataset | N/A | • Hill pars of h($[gTF]_i$)<br>• Hill pars of h($[PU]_i$) |
| C/EBP$\alpha$ model (without C/EBP$\alpha$ autoregulation) | • HSC differentiation dataset<br>• PU.1 transcriptional dataset | • $D_{CE}$<br>• $N_k$ of Biophysical($[PU]_i, [CE]_i$)<br>• $N$ of h($[PU]_i$) | • $\{G'^{\circ}\}$ of Biophysical($[PU]_i, [CE]_i$)<br>• Hill pars of h($[PU]_i$) except $N$<br>• $\varepsilon$ |
| C/EBP$\alpha$ model (with C/EBP$\alpha$ autoregulation) | • HSC differentiation dataset<br>• PU.1 transcriptional dataset | • $D_{CE}$<br>• $N_k$ of Biophysical($[PU]_i, [CE]_i$)<br>• $N$ of h($[PU]_i$)<br>• $N$ of h($[CE]_i$) | • $\{G'^{\circ}\}$ of Biophysical($[PU]_i, [CE]_i$)<br>• Hill pars of h($[PU]_i$) except $N$<br>• Hill pars of h($[CE]_i$) except $N$<br>• $\varepsilon$ |
| C/EBP$\alpha$ model (open-loop) | • HSC differentiation dataset<br>• PU.1 transcriptional dataset | • $N_k$ of Biophysical($[PU]_i, [CE]_i$) | • $\{G'^{\circ}\}$ of Biophysical($[PU]_i, [CE]_i$)<br>• $[CE]_i$ for all $i$<br>• $\varepsilon$ |
| RUNX1 model (without C/EBP$\alpha$ autoregulation) | • HSC differentiation dataset<br>• Expanded PU.1 transcriptional dataset | • $D_{CE}$<br>• $N_k$ of Biophysical($[PU]_i, [CE]_i, [RUN]_i$)<br>• $N$ of h($[PU]_i$) (it is later relaxed)<br>• $[RUN]_i$ | • $\{G'^{\circ}\}$ of Biophysical($[PU]_i, [CE]_i, [RUN]_i$)<br>• Hill pars of h($[PU]_i$) except $N$<br>• $\varepsilon$ |
| RUNX1 model (with C/EBP$\alpha$ autoregulation) | • HSC differentiation dataset<br>• Expanded PU.1 transcriptional dataset | • $D_{CE}$<br>• $N_k$ of Biophysical($[PU]_i, [CE]_i, [RUN]_i$)<br>• $N$ of h($[PU]_i$)<br>• $N$ of h($[CE]_i$)<br>• $[RUN]_i$ | • $\{G'^{\circ}\}$ of Biophysical($[PU]_i, [CE]_i, [RUN]_i$)<br>• Hill pars of h($[PU]_i$) except $N$<br>• Hill pars of h($[CE]_i$) except $N$<br>• $\varepsilon$ |
| RUNX1 model (with RUNX1 activation) | • HSC differentiation dataset<br>• Expanded PU.1 transcriptional dataset | • $D_{CE}$<br>• $N_k$ of Biophysical($[PU]_i, [CE]_i, [RUN]_i$)<br>• $N$ of h($[PU]_i$)<br>• $N$ of h($[RUN]_i$)<br>• $[RUN]_i$ | • $\{G'^{\circ}\}$ of Biophysical($[PU]_i, [CE]_i, [RUN]_i$)<br>• Hill pars of h($[PU]_i$) except $N$<br>• Hill pars of h($[RUN]_i$) except $N$<br>• $\varepsilon$ |

Table S8: Summary of experimental evidence that qualitatively or quantitatively constrains the parameters in the biophysical model.

| **Numerical constraint/Mechanism** | **Model component** | **Supporting experimental evidence** |
|---|---|---|
| Cooperativity of PU.1, C/EBP$\alpha$, and RUNX1 in the biophysical function Biophysical(...) is 1, 2, and 2, respectively | $N_k$ of Eqn. (S30). $N_k = 1$ if $[L_k] = [PU]$; $N_k = 2$ if $[L_k] = [CE]$ or $[RUN]$ | • Monomer form of PU.1 binds to DNA during cell differentiation [12]; PU.1 does not homodimerize to bind to DNA [13]<br>• C/EBP$\alpha$ binds to DNA as a dimer [14]<br>• RUNX1 binds to DNA as a dimer [15] |
| PU.1 and C/EBP$\alpha$ are in a positive feedback loop | Formulation of all models containing PU.1-C/EBP$\alpha$ positive feedback in Table S2 | • C/EBP$\alpha$ binds and regulates PU.1 promoter: Figures 2–4 of [16]<br>• C/EBP$\alpha$ binds to PU.1 upstream regulatory element: Figures 2 and 3 of [17]<br>• PU.1 binds to and regulates the +37 kb *Cebpa* enhancer: Figures 4 and 5D of [4]<br>• PU.1 and C/EBP$\alpha$ expression increases concurrently as HSCs differentiate from PU.1$^{\mathrm{lo}}$ MPP and PU.1$^{+}$ MPP (Lin$^{-}$Sca-1$^{+}$c-Kit$^{+}$CD34$^{+}$Flt3$^{+}$): Figures 2A and 6C of [18]<br>• *Cebpa* and *Spi1* expression increases from HSC1 (Lin$^{-}$CD133$^{+}$CD34$^{+}$) to MONO1 (CD34$^{-}$CD33$^{+}$CD13$^{+}$): Figure S6G and I of [19] |
| PU.1 activates C/EBP$\alpha$ with a cooperativity of 2 | $N$ in h($[PU]_i$) of $\frac{d[CE]_i}{dt}$ in Table S2 | • There are two PU.1 binding sites on the +37 kb *Cebpa* enhancer: Figure 4A, "ETSf" and "ETSb" of [4]<br>• Monomer form of PU.1 binds to DNA during cell differentiation [12]; PU.1 does not homodimerize to bind to DNA [13] |
| C/EBP$\alpha$ activates its own expression with cooperativity of 4 | $N$ in h($[CE]_i$) of $\frac{d[CE]_i}{dt}$ in Table S2 | • There are two C/EBP$\alpha$ binding sites on the +37 kb *Cebpa* enhancer: Figure 1 of [4]<br>• C/EBP$\alpha$ binds to DNA as a dimer [14] |
| RUNX1 activates C/EBP$\alpha$ with a cooperativity of 8 | $N$ in h($[RUN]_i$) of $\frac{d[CE]_i}{dt}$ in Table S2 | • There are four RUNX1 binding sites on the +37 kb *Cebpa* enhancer: Figure 1A of [4]<br>• RUNX1 binds to DNA as a dimer [15] |
| Transcription factors bound to the PU.1 enhancer destabilize the formation of nucleosomes | Nonzero $G^{\circ}_{kj}$, with $\sigma_k = 1^*$ and $\sigma_j = 1$, for $k \in \{1, 2, 3 \ldots T, \text{pro}\}$ and $j \in \{1, 2, 3 \ldots T\}$ | • PU.1 recruits SWI/SNF chromatin remodeling complexes to increase chromatin accessibility: Figure 1 of [20]<br>• C/EBP$\alpha$ interacts with and possibly recruits histone acetyltransferase p300: Figure 4 of [21]<br>• p300 binds to the upstream regulatory element of PU.1: Figure 4C of [3]<br>• C/EBP$\alpha$ expression increases PU.1 occupancy and H3K9ac: Figure 10A–C of [3]<br>• RUNX1 induction increases H3K9ac abundance: Figure 6D of [22] |
| A DNA loop is formed between the upstream regulatory element and the promoter of PU.1, possibly enabling the interaction between RNAP and transcription factors that bind to the PU.1 enhancer | Nonzero $G^{\circ}_{kj}$, with $\sigma_k = 1$ and $\sigma_j = 1$, for $k \in \{1, 2, 3 \ldots T\}$ and $j \in \{\text{pro}\}$ | • Binding of the nuclear adaptor protein LDB1 (required for chromosomal looping) requires PU.1: Figure 5D of [23]<br>• There is physical interaction between the upstream regulatory element (both 14 and 12 kilobases upstream) and the PU.1 promoter: Figures 6 and S6 of [24] |

*(continued on next page)*

| Numerical constraint/Mechanism | Model component | Supporting experimental evidence |
|---|---|---|
| There is a protein–protein interaction between every pair of TFs | Nonzero $G^{\circ}_{kj}$, with $\sigma_k = 1$ and $\sigma_j = 1$, for $k \in \{1, 2, 3 \dots T\}$ and $j \in \{1, 2, 3 \dots T\}$ | • PU.1 and C/EBP$\alpha$ work together to open closed chromatin: [25]<br>• AML1 and C/EBP$\alpha$ physically interact with each other: Figure 5 of [26]<br>• AML1 and PU.1 physically interact with each other: Figure 4 of [27]<br>• AML1 and C/EBP$\alpha$ cooperatively bind to DNA: Figure 3 of [27]<br>• AML1 and PU.1 can cooperatively activate gene expression: Figure 5 of [27] |
| C/EBP$\alpha$ half-life is ∼100 minutes | $D_{CE}$ | Figure 3h, line "−Ani" of [28] |
| RUNX1 half-life is ∼60 minutes | Not used directly. However, the short half-life suggests the RUNX1 steady state abundance can be approximated with the RUNX1 production rate. As such, experimental RNA-seq data for RUNX1 (which estimates RUNX1 production rate) constrain RUNX1 abundances in the model (see Section 5.2.3). | [11] |
| Ratios of C/EBP$\alpha$ production rates between cell types are known | Used for external validation after best fits | Haemopedia-Mouse-RNASeq for *Cebpa* (click here to view), group by "celltype": Multi Potential Progenitor, Macrophage Lineage, and B Cell Lineage [10] |
| Ratios of RUNX1 production rates (thus, RUNX1 abundances) between cell types are known | $[RUN]_i$ in Biophysical$([PU]_i, [CE]_i, [RUN]_i)$ in Table S2 | Haemopedia-Mouse-RNASeq for *Runx1* (click here to view), group by "celltype": Multi Potential Progenitor, Macrophage Lineage, and B Cell Lineage [10] |

### 6 Understanding model nullclines

This section includes a detailed analysis of the nullclines for the dimensionless "C/EBP$\alpha$ model (without C/EBP$\alpha$ autoregulation)" and "RUNX1 model (without C/EBP$\alpha$ autoregulation)".

The nullcline of PU.1 is obtained by setting $\frac{d[pu]_i}{dt} = 0$:

$$\frac{\text{Production}([pu]_i[PU^*]_M, Z_i)}{\text{Production}([PU^*]_M, Z_M^*)} = \frac{\frac{\log 2}{\tau_i} + D_{PU}}{\frac{\log 2}{\tau_M} + D_{PU}}[pu]_i \tag{S68}$$

or

$$[pu]_i = \frac{\text{Production}([pu]_i[PU^*]_M, Z_i)}{\text{Production}([PU^*]_M, Z_M^*)} \times \frac{1}{\frac{\frac{\log 2}{\tau_i}+D_{PU}}{\frac{\log 2}{\tau_M}+D_{PU}}} \tag{S69}$$

In both biophysical models, the production rate function Production(...) is a biophysical function that depends on both abundances of PU.1 and C/EBP$\alpha$. As such, according to Eqn. (S47),

$$[pu]_i = \frac{\text{Prob}_{\text{trc}}([pu]_i, [ce]_i, \{G'^{\circ}\})}{\text{Prob}_{\text{trc}}(1, 1, \{G'^{\circ}\})} \times \frac{1}{\frac{\frac{\log 2}{\tau_i}+D_{PU}}{\frac{\log 2}{\tau_M}+D_{PU}}} \tag{S70}$$

For the C/EBP$\alpha$ nullcline, or $\frac{d[ce]_i}{dt} = 0$, similarly, it is equivalent to plotting

$$[ce]_i = \frac{\text{h}([pu]_i, K')}{\text{h}(1, K')} \times \frac{1}{\frac{\frac{\log 2}{\tau_i}+D_{CE}}{\frac{\log 2}{\tau_M}+D_{CE}}} \tag{S71}$$

based on the C/EBP$\alpha$ model equations defined in Table S2 and the normalization operation in Eqn. (S42).

Eqns. (S70) and (S71) show that the model nullcline is equivalent to plotting the dimensionless production rate of each transcription factor, scaled by some cell-type-specific constant. Therefore, the change in nullclines from Figure 5E to 5F in the manuscript is due to a higher production rate at the same C/EBP$\alpha$ abundance.

#### 7 Supplementary Figures

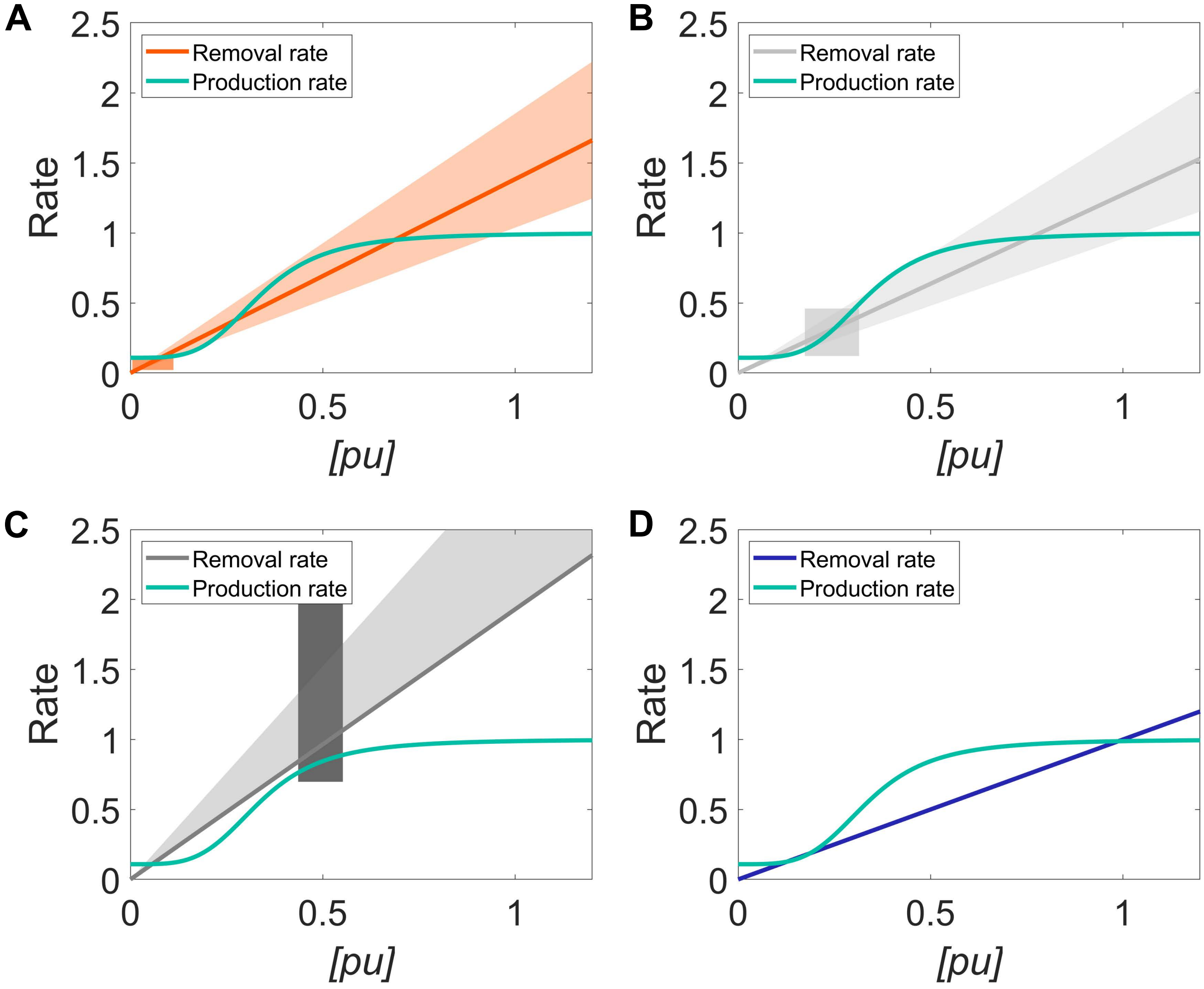

Figure S1: For the “AR&DGR model”, the fitted phenomenological production function is plotted against the protein removal function (constrained by cell-cycle length) of the **(A)** B cell, **(B)** pro-low, **(C)** pro-high, and **(D)** macrophage.

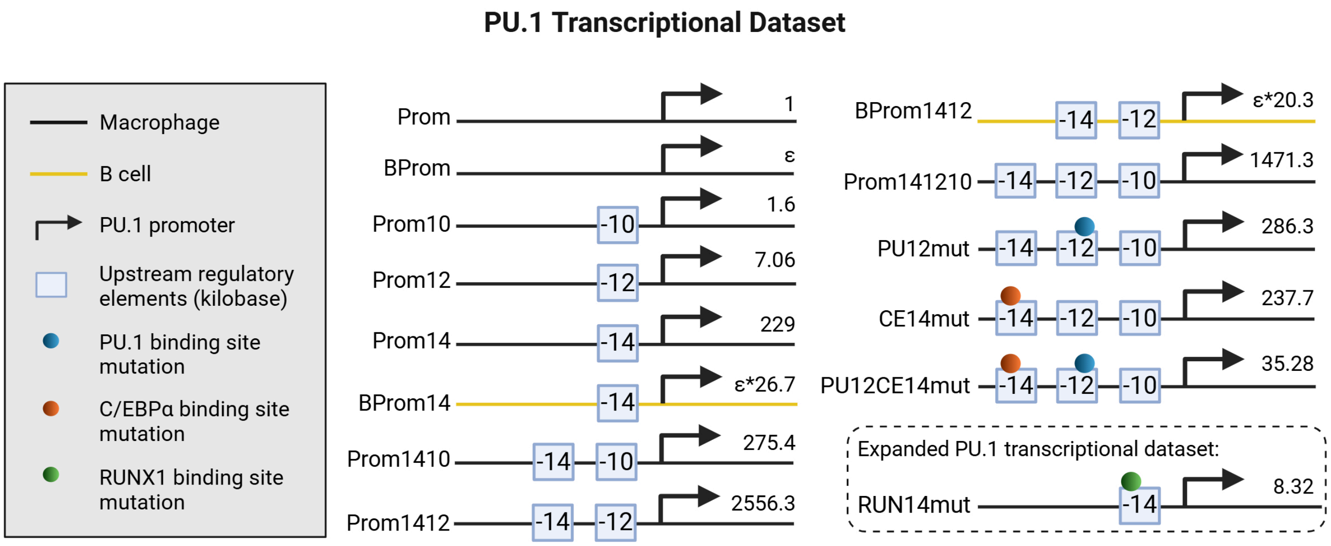

Figure S2: The complete list of gene constructs is illustrated graphically, along with their reporter activity measurements, which are used to constrain the biophysical models. "(Expanded) PU.1 transcription dataset" includes measurements from gene constructs without (with) "RUNX14mut".

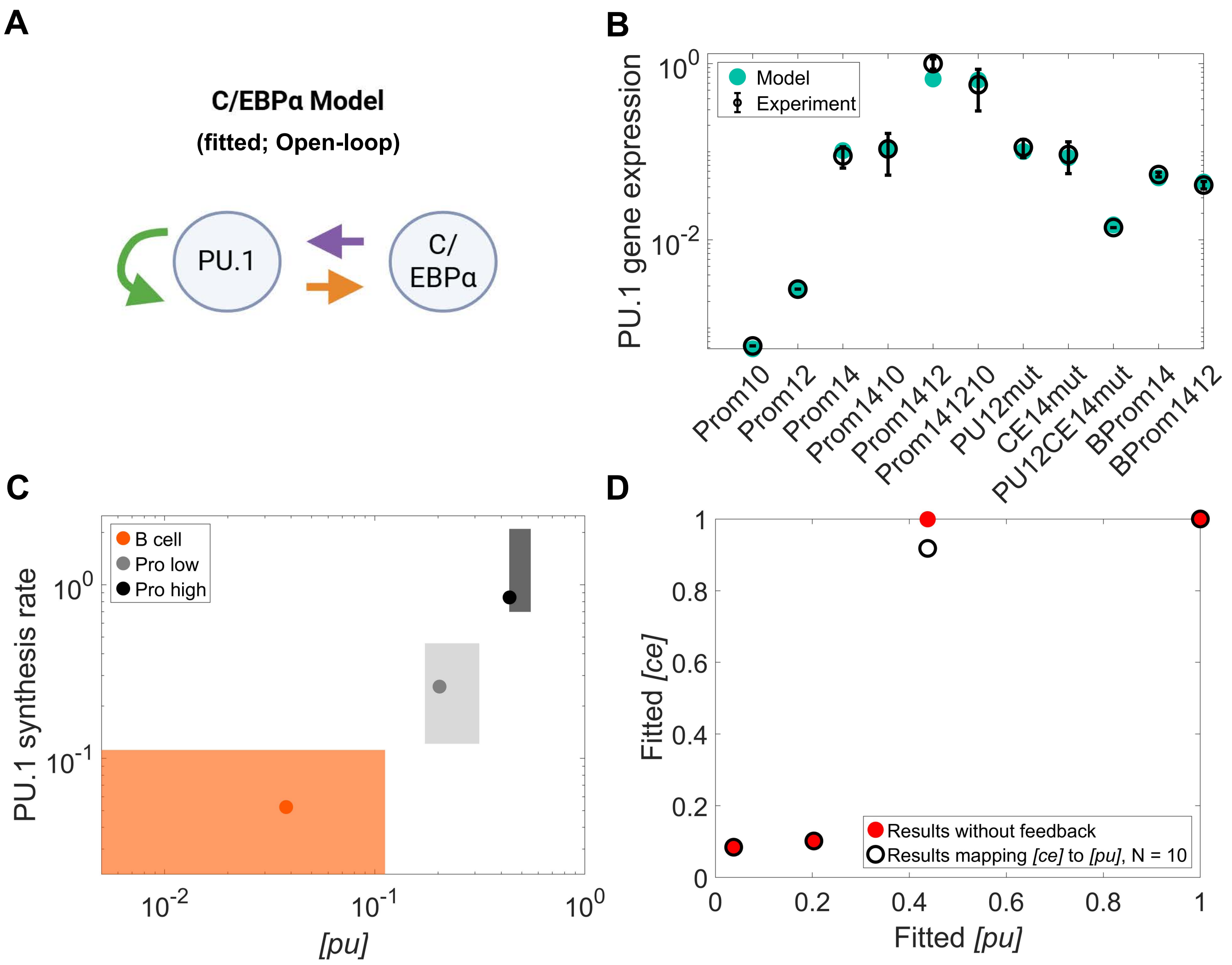

Figure S3: **(A)** The architecture of the open-loop "C/EBP$\alpha$ model" is shown. **(B)** Fitted reporter activities from the model in (A) are compared with the "PU.1 transcriptional dataset". **(C)** Fitted PU.1 abundances and synthesis rates from the model in (A) are compared with the joint measurement ranges. **(D)** Fitted C/EBP$\alpha$ and PU.1 abundance from the model in (A) were further fitted to an expression that is derived from the closed-loop model. This expression relates two transcription factor's abundance under PU.1-C/EBP$\alpha$ feedback (Supplementary Section 4). Compared to the PU.1-to-C/EBP$\alpha$ relationship found by the open-loop model (red filled circle markers), only a cooperativity of 10 in the PU.1-dependent C/EBP$\alpha$ activation leads to good fits. Given the PU.1 abundance found from the open-loop model, C/EBP$\alpha$ abundances fitted to the abovementioned closed-loop expression are plotted in black unfilled circle markers.

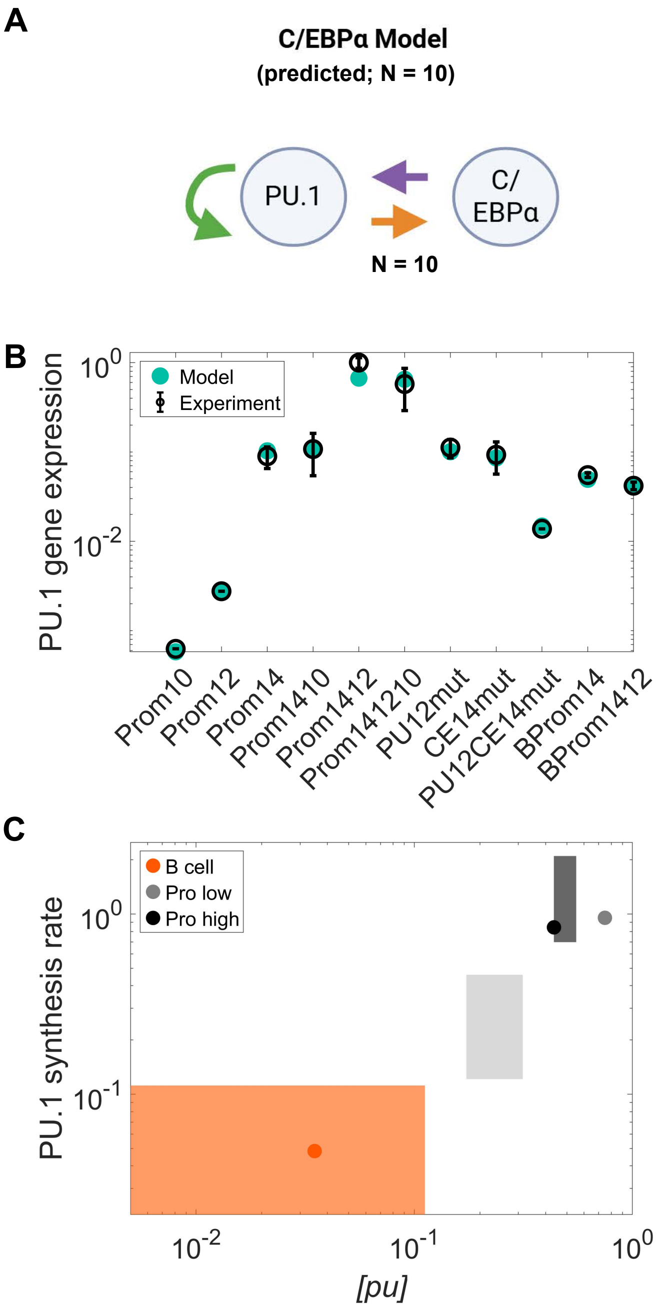

Figure S4: **(A)** The architecture of the closed-loop, high-cooperativity "C/EBP$\alpha$ model" is shown. This model was not fitted. Instead, we fed in best-fit open-loop parameters obtained from Figure S3B and C, combined with best-fit Hill parameters from Figure S3D, to examine whether an equivalent closed-loop model fits well. **(B)** Predicted reporter activities from the model in (A) are compared with the "PU.1 transcriptional dataset". **(C)** Predicted PU.1 abundances and synthesis rates from the model in (A) are compared with the joint measurement ranges.

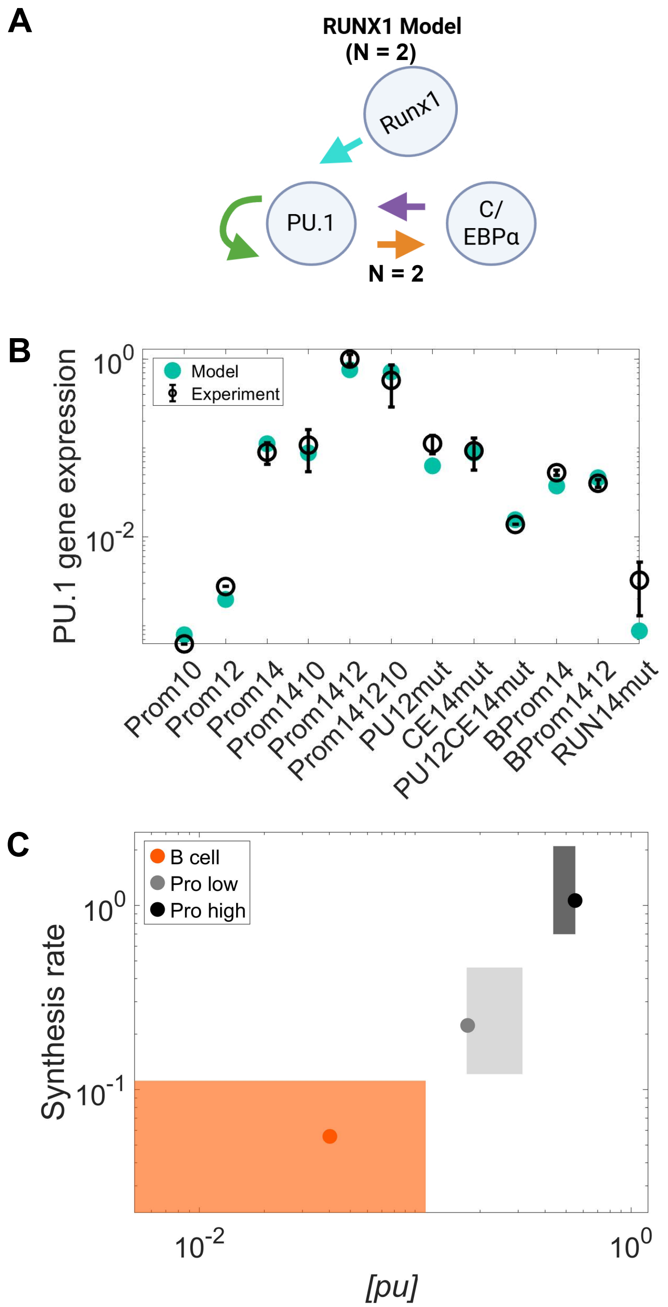

Figure S5: **(A)** The architecture of the low-cooperativity "RUNX1 model" is shown. **(B)** Fitted reporter activities from the model in (A) are compared with the "expanded PU.1 transcriptional dataset". **(C)** Fitted PU.1 abundances and synthesis rates from the model in (A) are compared with the joint measurement ranges.

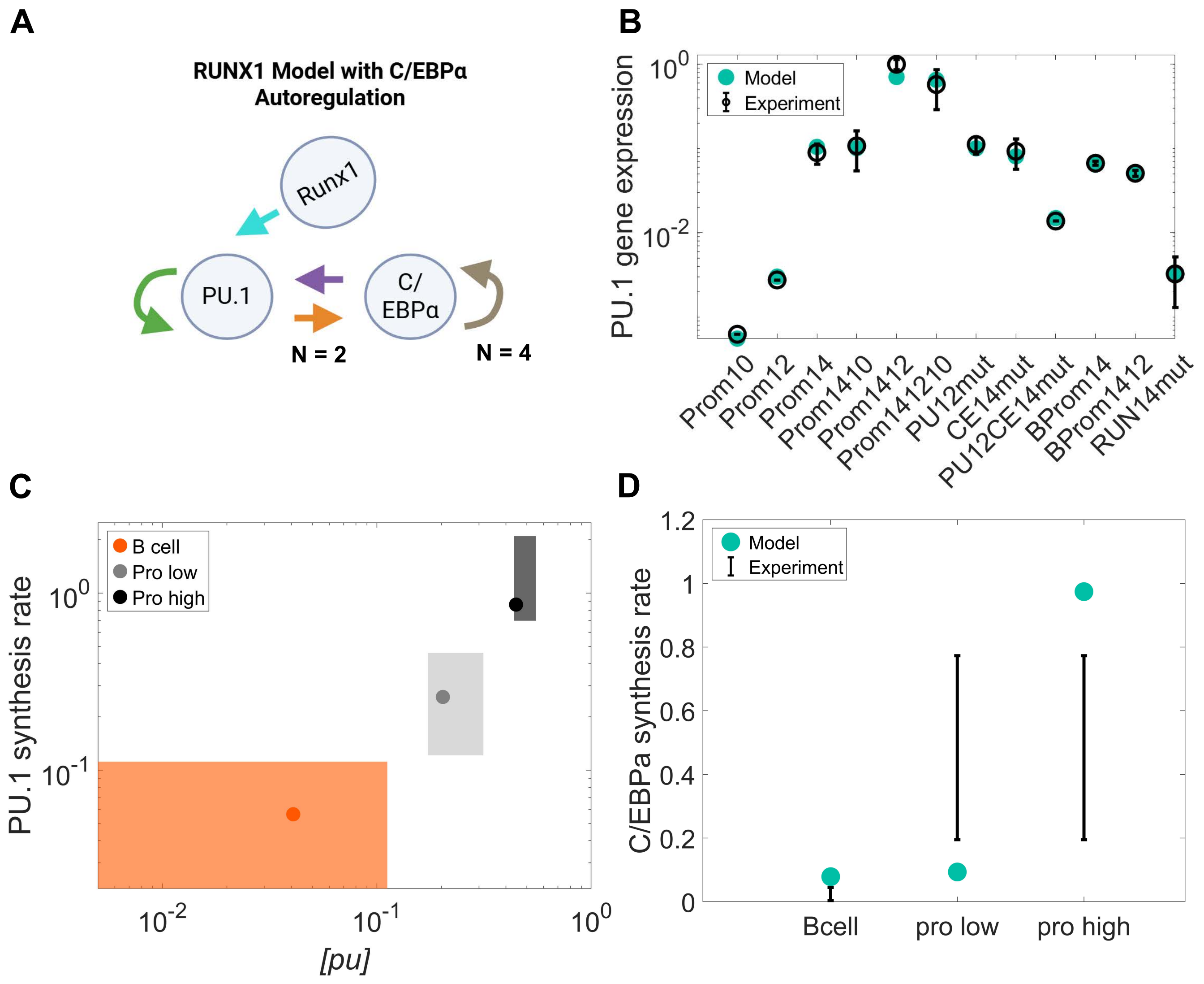

Figure S6: **(A)** The architecture of the low-cooperativity "RUNX1 model" with an additional C/EBP$\alpha$ autoregulatory feedback is shown. **(B)** Fitted reporter activities from the model in (A) are compared with the "expanded PU.1 transcriptional dataset". **(C)** Fitted PU.1 abundances and synthesis rates from the model in (A) are compared with the joint measurement ranges. **(D)** The C/EBP$\alpha$ production rates predicted by the model in (A) are compared with the experimental *Cebpa* RNA-seq measurements.

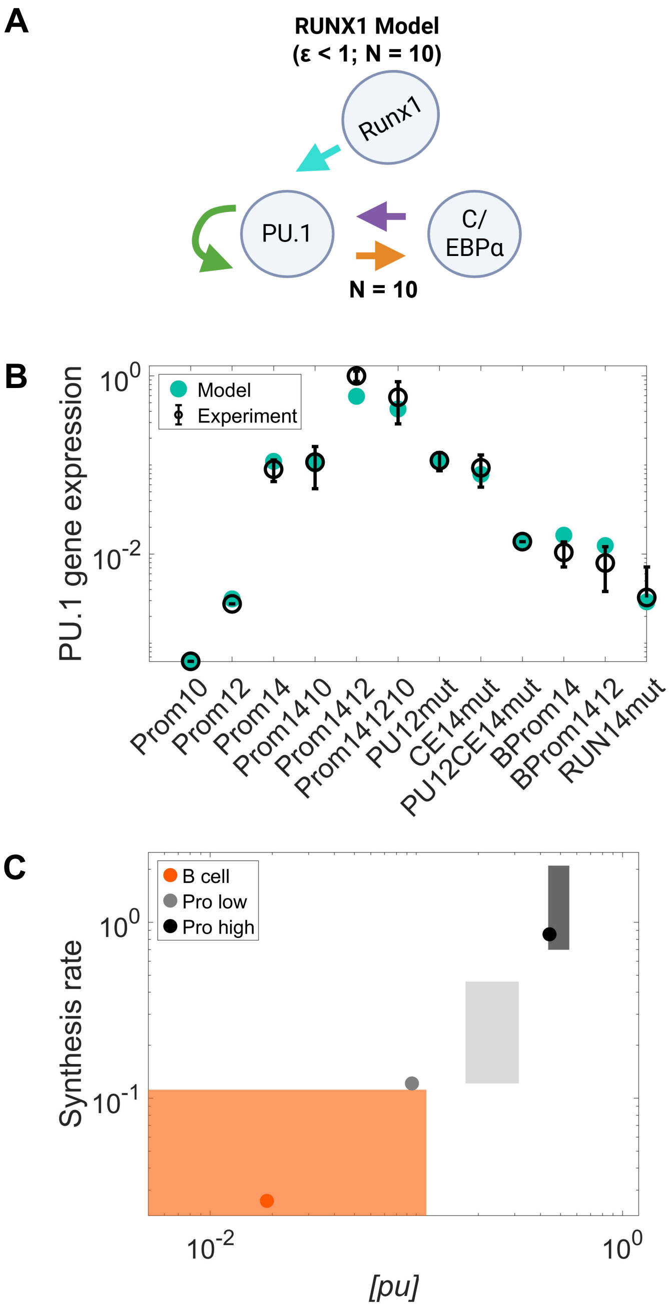

Figure S7: **(A)** The architecture of the high-cooperativity "RUNX1 model" is shown, where the B-cell-to-macrophage gene expression ratio ($\varepsilon$) is also constrained to be less than 1. **(B)** Fitted reporter activities from the model in (A) are compared with the "expanded PU.1 transcriptional dataset". **(C)** Fitted PU.1 abundances and synthesis rates from the model in (A) are compared with the joint measurement ranges.

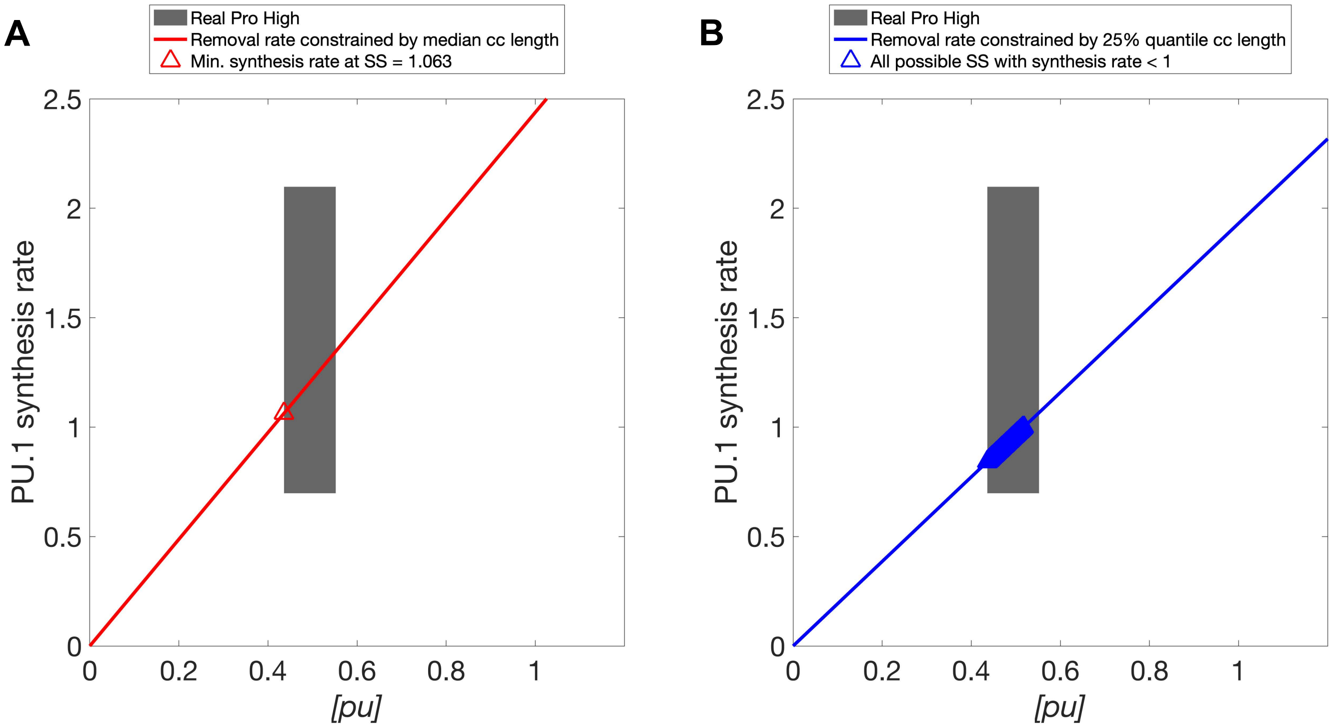

Figure S8: To find steady state ("SS") for pro-high systems, the comparison of using mean or 75$^{\text{th}}$ quantile of the measured cell cycle length ("cc length") as model constraints is shown. PU.1 removal rates across a testing range of PU.1 abundance (*[pu]*) are plotted in solid lines, computed from **(A)** the mean cc length and **(B)** the 75$^{\text{th}}$ quantile of cc length. In both plots, the joint measurements of PU.1 abundance and PU.1 synthesis rate with uncertainty is shown as a gray patch. At SS, the PU.1 production rate must equal to the PU.1 removal rate. Therefore, for any model to be considered successful, the production rate function must intersect at the region where solid lines (red in (A) and blue in (B)) overlaps with the gray patch. In (A), when removal rate is constrained by the mean cc length, the minimum ("min.") possible production rate at SS (triangle unfilled marker) is already greater than 1. However, with the tested model classes in our work, the production rate of pro-high cell state cannot exceed 1 (detailed explanation is provided in Supplementary Section 2.3). As such, we choose to use the 75$^{\text{th}}$ quantile as removal rate constraints, where, as shown in (B), the production rate is allowed to be smaller than 1 at various possible SS.

[9] Particle Swarm Optimization - MATLAB & Simulink. MathWorks, 2026.

[10] Jarny Choi, Tracey M Baldwin, Mae Wong, Jessica E Bolden, Kirsten A Fairfax, Erin C Lucas, Rebecca Cole, Christine Biben, Clare Morgan, Kerry A Ramsay, Ashley P Ng, Maria Kauppi, Lynn M Corcoran, Wei Shi, Nick Wilson, Michael J Wilson, Warren S Alexander, Douglas J Hilton, and Carolyn A de Graaf. Haemopedia RNA-seq: A database of gene expression during haematopoiesis in mice and humans. *Nucleic Acids Research*, 47(D1):D780–D785, January 2019.
